# H3.3 De Novo Mutations Alter Lysine 36 Methylation via Distinct Mechanisms

**DOI:** 10.1101/2025.09.09.674984

**Authors:** Shahir M. Morcos, Sabina Sarvan, Alejandro Saettone, Cassandra J. Wong, Samuel Chau Duy Tam Vo, Ayala Milo, Omar H. Bayoumy, Rajesh Angireddy, Giovanni L. Burke, Jack F. Greenblatt, Nada Jabado, Carol C.L. Chen, Elizabeth J. Bhoj, Anne-Claude Gingras, Jean-François Couture, Eric I. Campos

**Affiliations:** Genetics & Genome Biology Program, The Hospital for Sick Children, Toronto, Canada; Department of Molecular Genetics, University of Toronto, Toronto, Canada; Ottawa Institute of Systems Biology, Department of Biochemistry, Microbiology and Immunology, Faculty of Medicine, University of Ottawa, Ottawa, Canada; Lunenfeld-Tanenbaum Research Institute, Mount Sinai Hospital, Sinai Health System, Toronto, Canada; Center for Applied Genomics, Children’s Hospital of Philadelphia, Philadelphia, USA; Donnelly Centre, University of Toronto, Toronto, M5S 3E1, Canada; Department of Human Genetics, McGill University, Montreal, Canada; Department of Pediatrics, McGill University, and The Research Institute of the McGill University Health Centre, Montreal, Canada; Division of Experimental Medicine, Department of Medicine, McGill University, Montreal, Canada; Terry Fox Laboratory, BC Cancer, Vancouver, Canada; Department of Biochemistry and Molecular Biology, University of British Columbia, Vancouver, Canada

## Abstract

Bryant-Li-Bhoj syndrome (BLBS) is caused by de novo mutations on histone H3.3 and is generally characterized by severe neurodevelopmental deficits. Oncogenic H3.3 amino acid substitutions were described over the past decade, but the molecular impact of BLBS mutations remained unstudied. The remarkable number and spread of the missense mutations led us to hypothesize that some converge on the same downstream effectors.

We recently showed that H3.3G34R/V substitutions, seen in both cancer and BLBS, impair associations with the DNMT3A DNA methyltransferase. Our proteomic, enzymatic, and structural analyses now show that H3.3 BLBS mutations flanking glycine 34 have surprisingly stark effects on H3K36 methyltransferases, drastically altering H3K36 methyl states in cis and the binding of effector proteins with a PWWP domain, including DNMT3A/B. That confirms the existence of molecular commonalities amongst BLBS H3.3 point mutants while providing some of the first mechanistic insights into the syndrome.

## Introduction

Histone variants, their post-translational modifications (PTMs), and the proteins that bind them locally influence DNA fiber accessibility and transcriptional outputs^1,2^. Mammalian histone H3.1 is highly expressed in S-phase and is the predominant H3 variant in rapidly cycling somatic cells^3-5^. The H3.3 variant is instead basally expressed through the cell cycle and deposited in a DNA synthesis-independent manner, and accumulates in post-mitotic cells^3,5,6^. The HIRA histone chaperone complex generally deposits the histone on transcribed genes, while ATRX-DAXX deposits it on repetitive DNA elements^7-9^.

H3.3 is expressed from two genes, *H3-3A* and *H3-3B*, each having a unique promoter and expression pattern^10,11^. Murine *H3f3a* and *H3f3b* double knockouts result in early embryonic lethality^12,13^. Mice with both genes knocked out in neural progenitor cells also die shortly after birth, with negative impacts seen on neuronal fate and development^13,14^.

Histone genes are highly preserved amongst eukaryotes and pathogenic H3.3 K27M and G34R/V amino acid substitutions, first identified in pediatric brain tumors, grossly deregulate the epigenome^15-18^. Protein modeling showed that the bulkier side chains on G34 mutants create steric clashes with the catalytic SET domain of H3 lysine 36 methyltransferases^19,20^. The result is a loss in cis of the H3 lysine 36 di- and tri-methyl marks (H3K36me2/3), respectively catalyzed by NSD1/2 and SETD2.

SETD2 interacts and travels with the elongating Pol II machinery, trimethylating H3K36 on transcribed genes^21^ while NSD1/2 primarily install H3K36me2 over large intergenic regions^22,23^. These marks recruit proteins with a Proline-Tryptophan-Tryptophan-Proline (PWWP) domain, including the DNMT3A and DNMT3B de novo DNA methyltransferases that preferentially bind H3K36me2 and H3K36me3, respectively^24,25^. DNMT3A installs CG and non-CG DNA methyl marks — the latter being particularly important for stem cell differentiation and synaptogenesis^26^.

Over 60 distinct *H3-3A* and *H3-3B de novo* dominant missense mutations were identified in 98 patients with the recently described Bryant-Li-Bhoj neurodevelopmental syndrome (BLBS)^27-33^. A striking quarter or so of all H3.3 residues are concerned, spanning nearly the entire length of the protein. While phenotypic variability is observed amongst patients, the commonly shared traits, such as severe intellectual disability or a failure to reach important milestones (such as sitting, speaking, or walking), also imply molecular commonalities.

BLBS point mutants include the H3.3G34R/V substitutions. Mice with G34R directly knocked into the *H3f3a* gene exhibit fully penetrant microcephaly and progressive neurodegeneration, while the G34V substitution caused milder phenotypes^30^. At the cellular level, a depletion of intergenic H3K36me2 and DNMT3A, and a redistribution of the DNA methyltransferase over CpG islands caused pathogenic transcriptional changes. Because histones recruit effector proteins to chromatin, we sought to identify changes in protein associations amongst H3.3 BLBS point mutations flanking the H3.3G34 residue.

Distinct molecular mechanisms converged on SETD2 and NSD1/2 deregulation, thereby altering H3.3 associations with proteins that have a PWWP domain. While amino acid substitutions on the histone tail impact the catalytic SET domain of the enzymes, substitutions on the H3.3 N-terminal α-helix instead affected their activity via their Associated with SET (AWS) domain. Our DNMT3B PWWP crystal structure corroborates a model where the altered PWWP protein recruitment is largely due to the changes in H3.3K36me2/3 levels.

## Results

### Proteomic Analysis of BLBS Mutants Flanking H3.3 Glycine 34

To define the impact of BLBS mutations on H3.3 protein interactions, we generated a library of isogenic cells expressing WT H3.3 or one of 10 missense substitutions (7 flanking G34, and 3 at a distal location – **Fig. 1a**). Proximal protein associations were identified using BioID^34^ in the inducible HEK293 Flp-In T-REx cell model, previously used to assess histone proteomes^30,35-37^. Each lineage expressed the histone constructs at comparable levels from a single defined genomic locus. The BioID2 enzyme^38^ was fused to either of the N- and C-terminal H3.3 extremities, and each BioID experiment was done in duplicates for a total of four analyses per histone (**Fig. 1b**). The exogenous histones incorporated chromatin and represented less than 20% of the total H3.3 (**Suppl. Fig. 1**).

**Figure 1.**
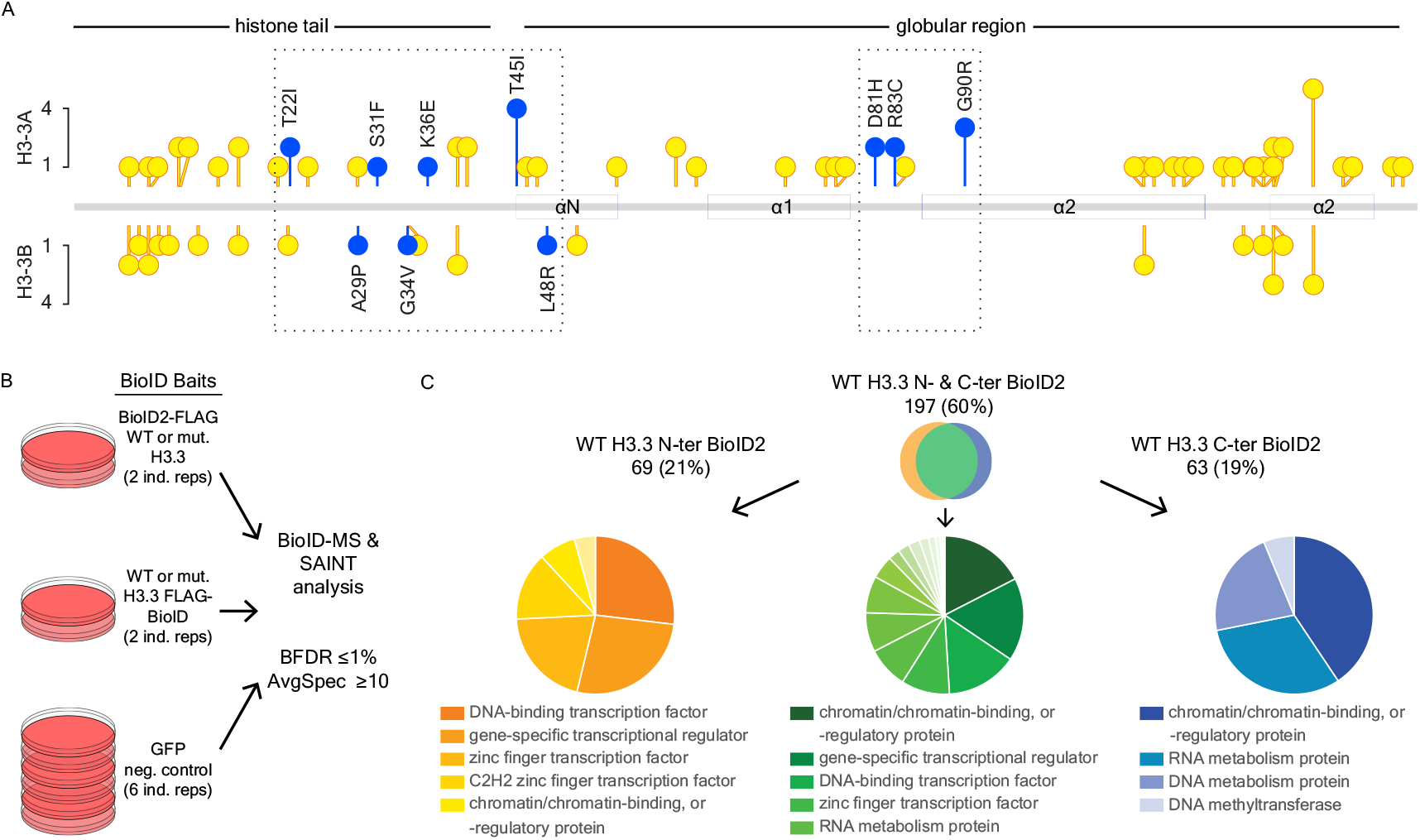
BioID analysis of H3.3 and BLBS missense mutations. **(a)** Distribution of BLBS amino acid substitutions on the histone. **(b)** BioID experimental pipeline. **(c)** Venn diagram showing the overlap between proximal associations identified for WT H3.3 with BioID2 fused at the N- and C-termini. Pie charts provide an overview of the commonly represented PANTHER protein classes, of which the top-most are labeled^64,65^.

The WT H3.3 BioID2 constructs together identified over 350 proximal associations, with nearly two-thirds identified by both H3.3 N- and C-ter BioID2 fusions (**Fig. 1c**). For stringency, we only considered preys identified with ≥10 average spectral counts and a clear enrichment over the GFP-BioID2 control (BFDR ≤ 1%). There was a good agreement (60-63% overlap) with H3.3 associations identified using a different biotin ligase^35^ and with those annotated in the BioGrid database^39^. Nearly 85% of the mutant H3.3 proximal associations had average spectral values that deviated by <1.5-fold from those seen with WT H3.3. For simplicity, we hence refer to protein associations with ≥1.5-fold change as altered or deregulated.

We purposely included the BLBS H3.3 G90R mutant in our BioID screen since the glycine is unique to H3.3 and amino acid substitutions at this position abolish interactions with the H3.3-specific histone chaperone DAXX^40^. As expected, there was a complete loss of associations between H3.3 G90R and DAXX regardless of the BioID2 position on the histone (**Fig. 2a**), though DAXX fell shy of our BFDR and spectral count cutoffs with the H3.3 G90R N-terminal BioID2. The same was observed after immunoprecipitating the proteins (**Fig. 2B**). The notable decrease in ATRX associations but not a complete loss was again expected, since ATRX independently binds H3K9me3/H3K4me0 on chromatin via its ADD domain^41-43^.

**Figure 2.**
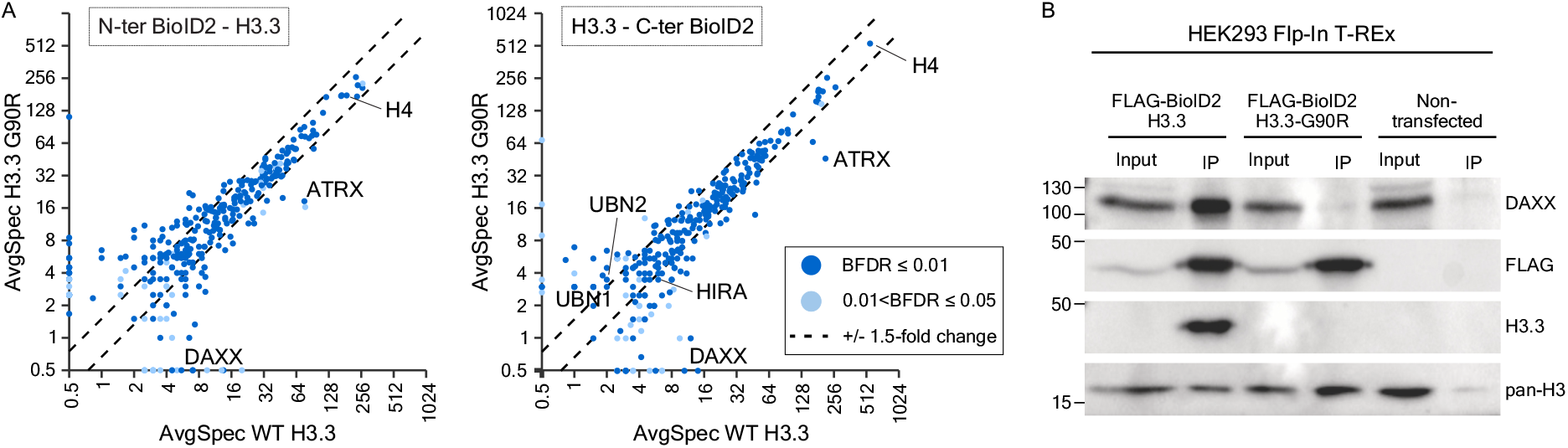
The H3.3G90R BLBS point mutant abrogates binding to the DAXX histone chaperone. **(a)** Scatter plots comparing average spectral counts (AvgSpec) of BioID preys identified by WT H3.3 (x axes) and the H3.3G90R (y axes) histone baits. Left and right panels respectively show results obtained with the N- and C-ter H3.3 fusions to BioID2. Preys were shown if enriched over a GFP BioID control (BFDR ≤5%). **(b)** Immunoprecipitation of N-ter tagged WT and G90R H3.3 showing the inability to coprecipitate DAXX with the N-ter BioID-FLAG-H3.3G90R construct. Inputs represented 5% of the sample material. Note that the H3.3-specific antibody binds the _87_AAIG_90_ H3.3 sequence and therefore does not recognize the H3.3G90R mutant.

The BioID revealed uniquely altered protein associations for each point mutant (**Suppl. Figs. 2-4**), but commonalities were also discernible. Eighty-five protein associations were similarly altered in at least 3 of the 10 H3.3 point mutants (**Suppl. Fig. 5**). The vast majority involved “chromatin-binding/-regulatory proteins” and transcription-related PANTHER protein class terms (**Fig. 3a**). Interestingly, prominent and opposite changes were observed for associations between the DNMT3A/B DNA methyltransferases and H3.3 point mutants other than G34V and K36E (**Fig. 3b, Suppl. Fig. 2-4**).

**Figure 3.**
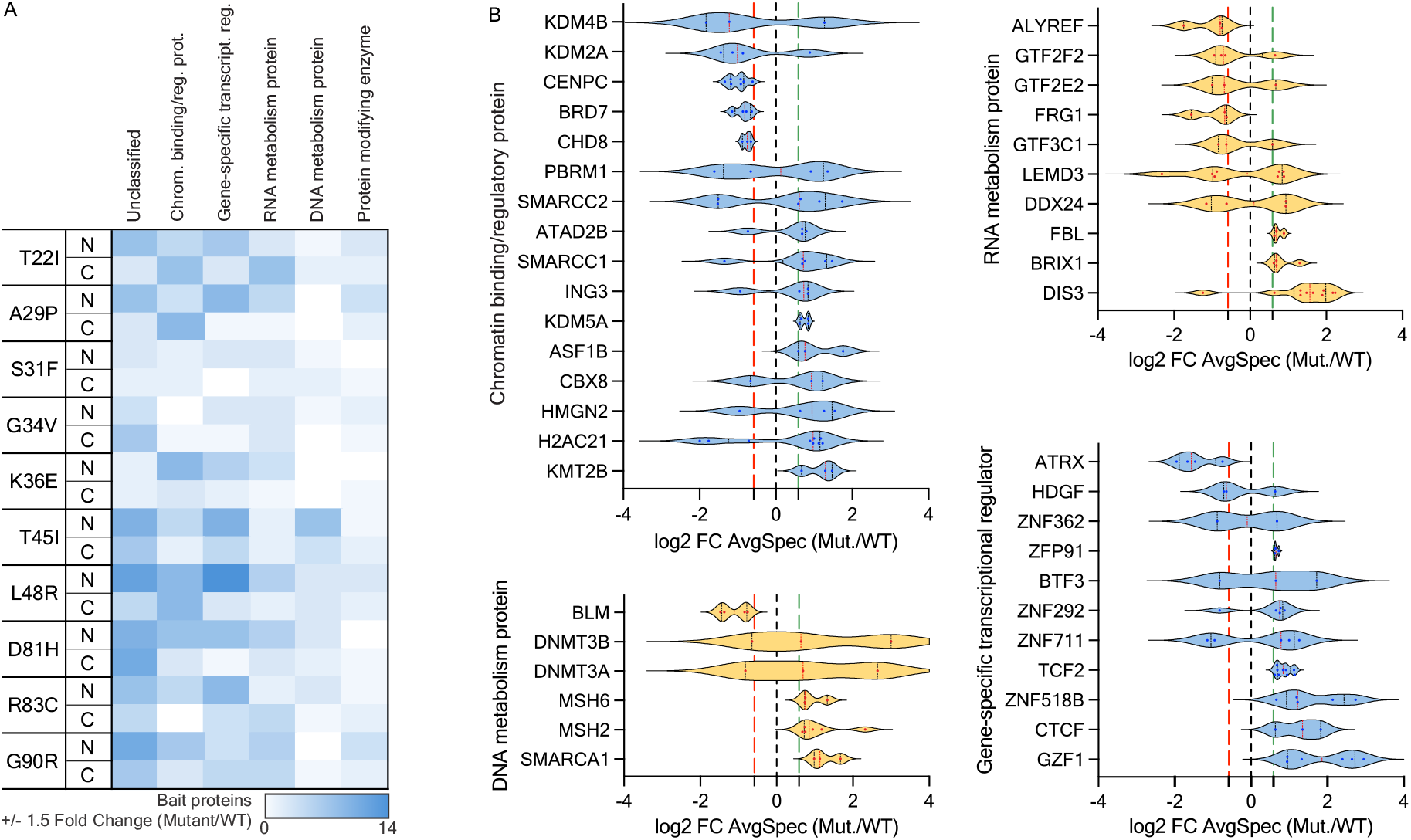
BLBS H3.3 point mutants have altered associations with chromatin-regulating proteins. **(a)** Heatmap showing the number of deregulated prey per PANTHER database^64,65^ protein class within both N- and C-ter baits. Protein classes with ≥ 4 deregulated prey in at least one mutation are shown. **(b)** Violin plots showing the distribution of mutant/WT average spectral counts amongst the most frequently deregulated protein classes. The red (leftmost) and green (rightmost) dashed lines mark a 1.5-fold change.

### H3.3 Point Mutants with Recurrently Altered PWWP Protein Associations

We next sought to identify patterns amongst commonly deregulated protein associations by grouping them based on function or the presence of shared histone-binding domains (**Fig. 4a, Suppl Fig. 6**). Interesting patterns emerged, with the most N-terminal point mutants in our cohort (T22I, A29P) gaining associations with bromodomain-containing proteins, for example. Proteins with a PWWP domain were among the most frequently disrupted groups, with significant changes for S31F, T45I, and L48R, and notably decreased associations between G34V, G90R and various PWWP proteins (**Fig. 4a-b, Suppl. Fig. 7**). The variability between the various H3.3-PWWP domain associations is worth mentioning. Associations with PSIP1, for example, were hardly affected by the point mutants, while those with BRD1, BRPF1 (peregrin), DNMT3A/B, MSH6, and ZMYND8 were frequently deregulated in the H3.3 S31F, G34V, K36E, T45I, L48R, and G90R point mutants (**Fig. 4b, Suppl. Fig. 7**). This can be attributed to intrinsic variability across PWWP domains and the presence of other protein-binding domains or motifs on these proteins and/or their binding partners. Interestingly, the associations between PWWP and the S31 and T45I point mutants increased while those with G34V, K36E, and L48R generally decreased.

**Figure 4.**
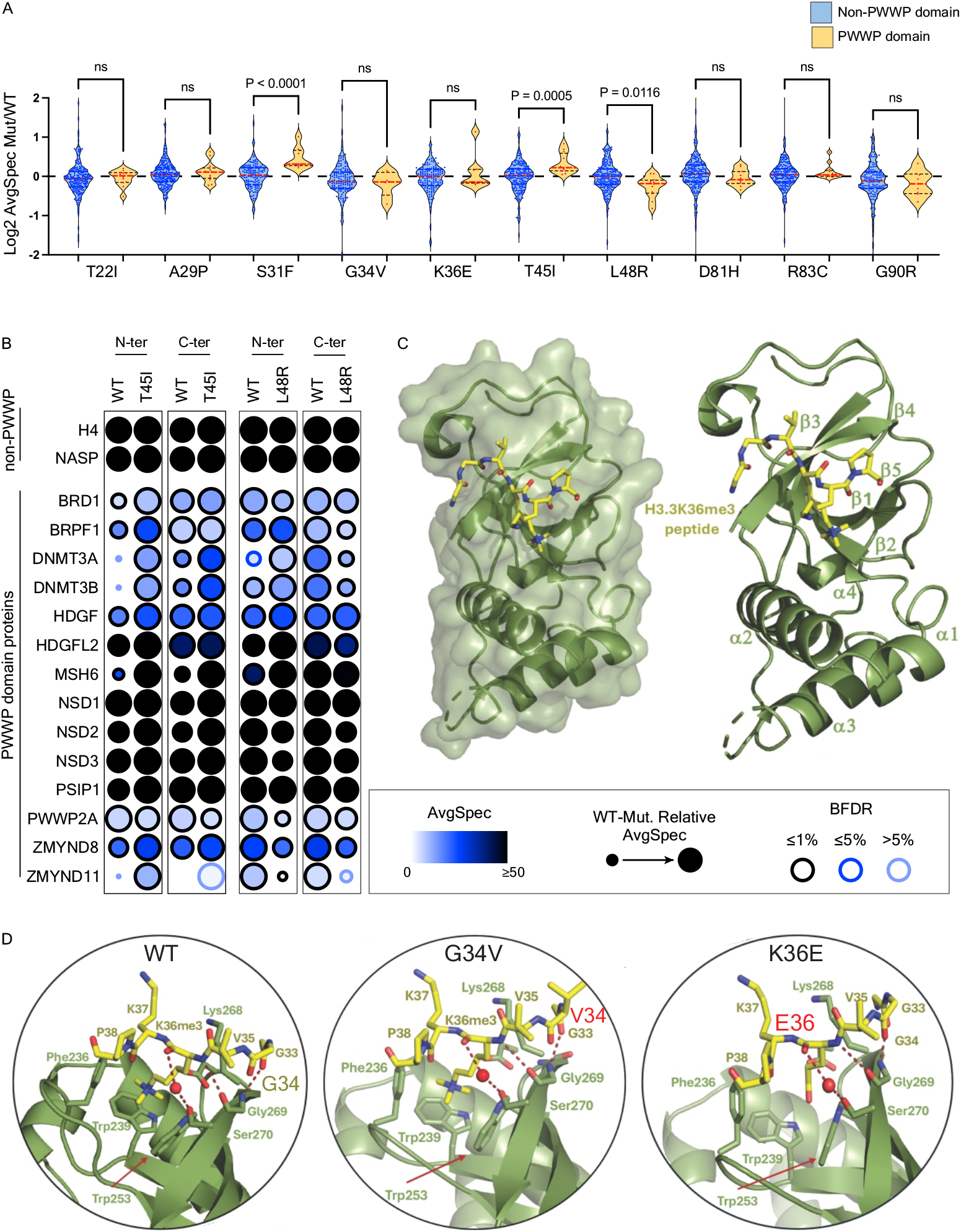
Altered associations between PWWP-containing proteins and missense mutants flanking H3.3K36. **(a)** Distribution of mutant/WT AvgSpec ratio for prey proteins with or without a PWWP protein domain, using the C-ter BioID2 bait proteins. Results of a Mann-Whitney statistical analysis are shown. **(b)** Dot plot showing prey proteins with a PWWP domain identified with the H3.3 T45I and L48R prey. **(c)** Cartoon representation of the crystal structure of the complex between the DNMT3B PWWP domain and H3K36me3 peptide (a.a. 21-46). Positively charged surfaces are colored in blue (+70 kbTe-1) and negatively charged surfaces in red (−70 kbTe-1) (kb = Boltzmann’s constant, T = temperature in Kelvin, and e = charge of an electron). Secondary structures of the DNMT3B PWWP domain and the histone residues are labelled. The leftmost image includes a surface representation of the DNMT3B PWWP domain, with the histone H3.3 peptide shown in stick. **(d)** Intermolecular interactions of the histone peptide with the PWWP domain. DNMT3B PWWP residues that form the histone H3.3 binding cleft are shown in stick representation and labelled. Red spheres represent water molecules and hydrogen bonds and are rendered in firebrick dashed lines. Predicted intermolecular interactions of histone H3.3 G34V and K36E are also shown.

### Structural Analysis of PWWP Associations with H3.3 BLBS Point Mutants

PWWP domains bind both H3K36me and nucleosomal DNA. Available structures suggest a short footprint for the BRPF1^44,45^, DNMT3B^46^, LEDGF^47,48^, and ZMYND11^49,50^ PWWP domains on the H3 tail. The LEDGF structures included whole nucleosome core particles, yet its interactions with H3 were again limited (mostly engaging residues between valine 35 and tyrosine 41).

DNMT3A/B were particularly affected by the BLBS H3.3 point mutants, so we next obtained the crystal structure of the DNMT3B PWWP in association with an H3 peptide (**Fig. 4c, Table 1**). While not the first such structure, the peptide used here was longer than the previously used one (a.a. 21-46 instead of 30-40). Akin to prior structures, the binding was restricted to a few histone residues (a.a. 33-38). Hence, the changes in the PWWP binding to H3.3 T45I and L48R could not be direct. Furthermore, the modelling of H3.3 G34V and K36E on that structure predicts little impact on the PWWP binding to the H3.3 tail (**Fig. 4d**).

**Table 1.** Data collection and refinement statistics for DNMT3B^PWWP^/H3.3 complex.

|  |  |
| --- | --- |
| **Data Collection** |  |
| Space group | P 3<sub>1</sub> 2 1 |
| Cell dimensions |  |
| a, b, c (Å) | 74.23, 74.23, 157.15 |
| α, β, γ (°) | 90, 90, 120 |
| Resolution | 20.6 - 2.4 (2.5 - 2.4) * |
| *R*<sub>meas</sub> | 0.05 (0.48) |
| *I* / σ*I* | 24.8 (5.8) |
| Completeness (%) | 99.8 (100.0) |
| Redundancy | 19.7 (19.6) |
| **Refinement** |  |
| Resolution (Å) | 20.6 – 2.4 |
| No. reflections | 20181 |
| *R*<sub>work</sub> / *R*<sub>free</sub> | 0.24 / 0.27 |
| Num. atoms |  |
| PWWP | 2076 |
| H3.3 | 73 |
| Water | 34 |
| *B*-factors (Å<sup>2</sup>) |  |
| PWWP | 33.3 |
| H3.3 | 31.2 |
| Water | 55.0 |
| R.m.s. deviations |  |
| Bond lengths (Å) | 0.002 |
| Bond angles (°) | 0.59 |
| Molprobity score | 1.69 |
| Clashscore | 5.4 |
| Ramachandran favored (%) | 96% |
| Ramachandran allowed (%) | 4% |
| * Highest-resolution shell is shown in parentheses. |  |

### H3.3 BLBS Point Mutants Affect H3K36 Methylation via Varied Molecular Means

The characteristic PWWP binding to methylated H3K36 hinted towards an upstream deregulation of the mark. Histones were hence acid-extracted from the isogenic cell lines expressing the C-terminal BioID2 fusion proteins. As expected, the western analysis for H3K36me3 generally followed the trends seen with PWWP proteins by BioID, with a bimodal signal distribution for A29P, increased H3.3K36me3 levels with S31F and T45I, and a complete to near complete loss of signal with G34V, K36E, and L48R (**Fig. 5a**). All the altered methyl levels occurred in cis and did not impact overall K36me on the endogenous histones. The increased methyl levels on the T45I mutant were particularly striking and were again seen after immunoprecipitating the N-ter tagged mutant (**Suppl. Fig. 8a**).

**Figure 5.**
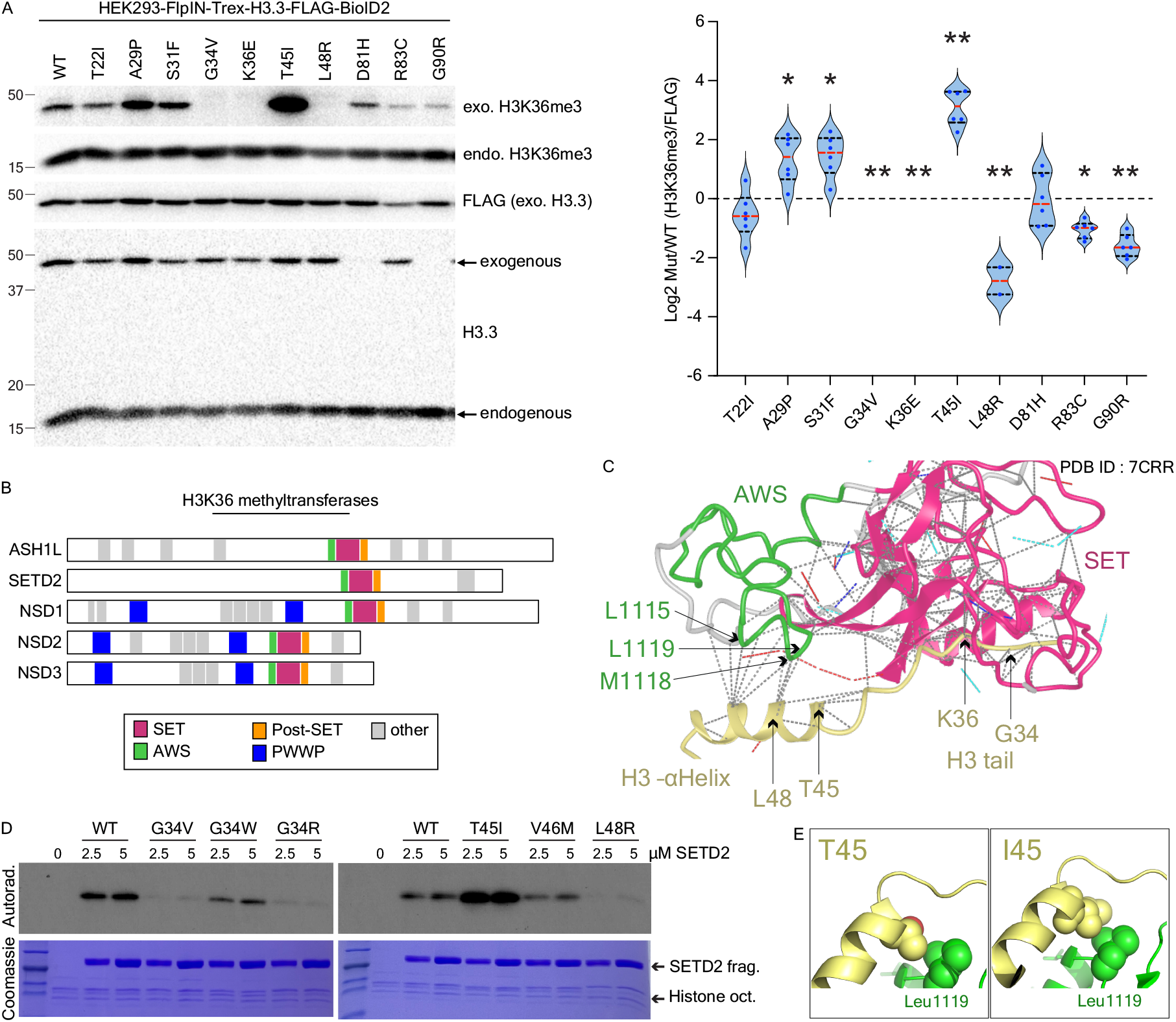
Altered H3K36 tri-methylation across various BLBS H3.3 mutants. **(a)** Western blot analysis of acid-extracted histones (n=6; 3 biological replicates, 2 technical replicates each). Image Lab was used to quantify the western signals. H3K36me3 signals were normalized to the total exogenous H3.3 (FLAG) and compared to the WT histone. Results of a Mann-Whitney statistical analysis are shown. No H3K36me3 signals were detected with G34V or K36E. **(b)** Primary structure of ASH1L, NSD1–3, and SETD2. **(c)** Zoomed view of the cryo electron microscopy structure of mammalian NSD3 in complex with a nucleosome (PDB: 7CRR)^51^, showing the interaction of the SET and AWS domain with histone H3. **(d)** In vitro methyltransferase assay using a SETD2 catalytic fragment composed of the AWS, SET, and Post-SET regions on recombinant nucleosome particles (one of 3-4 repetitions is shown; the full figure and quantification are available in **Suppl. Fig. 8b**). **(e)** Zoomed view of the T45 surface binding region of Set2. Mutation followed by energy minimization shows additional hydrophobic contacts between Ile45 and Leu1119.

The loss of H3.3K36 methylation on the G34V point mutant is attributed to steric clashes within the catalytic SET domain of H3K36 methyltransferases^19,20^. Interestingly, the altered H3.3K36 methyl levels on other BLBS point mutants were not caused by the same mechanism. The SETD2, NSD1-3, and ASH1L enzymes share the catalytic SET domain and a post-SET domain, but also uniquely possess an Associated With Set (AWS) domain only found on these HK36 methyltransferases (**Fig. 5b**). The crystal structure of NSD3 bound to a nucleosome core particle^51^ revealed extensive contacts between the AWS and the N-terminal α-helix of histone H3 (**Fig. 5c**). These notably involved hydrogen bonds between H3 R49/R52 and AWS L1116/L1119, further stabilized by Y1121 through hydrophobic stacking that, when disrupted, dramatically reduced the activity of the enzyme.

The T45I and L48R point mutants caused stark and opposite changes in PWWP associations (**Figs. 4a-b**) and H3.3K36me3 levels (**Fig. 5a**). This was intriguing given that the residues are only separated by 2 amino acids. In vitro enzymatic assays were performed to verify if these point mutants directly impact the catalytic activity of H3K36 methyltransferases.

Recombinant nucleosome core particles carrying either WT H3.3, G34R, G34V, G34W, T45I, V46M, or L48R were methylated using SETD2 (**Fig. 5d**) or NSD1 (**Suppl. Fig. 8**) catalytic fragments composed of the AWS and SET domains. G34R and G34W had the strongest and weakest impact on K36 methylation, respectively. Their impact on the SETD2 and NSD1 enzymatic activities was also in line with recently published data showing drastic epigenomic and phenotypic effects with the arginine substitution and the weakest with the tryptophan^30^.

Importantly, the in vitro methylation assays showed that H3.3 T45I stimulates the enzymatic activities of SETD2 and NSD1, and L48R abolishes it (**Fig. 5d, Suppl. Fig. 8**). Our modeling points to changes in hydrophobic interactions as a plausible molecular cause of the changes seen with T45I and L48R. Valine 46 does not contact SETD2, likely explaining why the V46M mutant had little impact on the SETD2 and NSD1 activities. In any case, it shows that the BLBS point mutants that impact H3.3K36me and PWWP binding do so through different means (**Fig. 6**).

**Figure 6.**
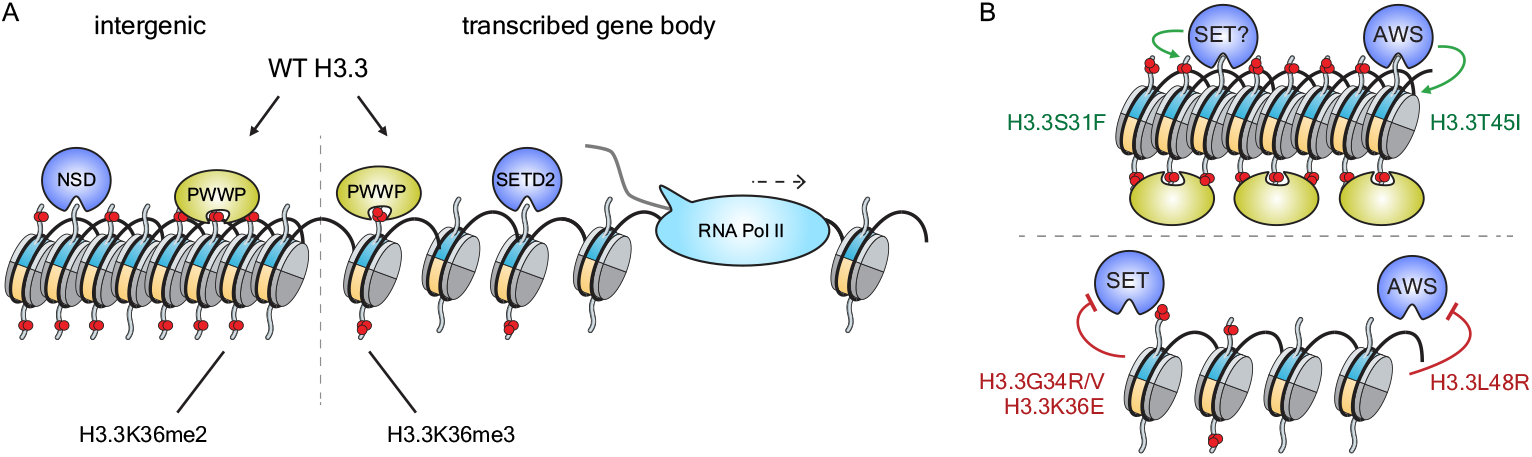
Model for H3K36me deregulation by a subset of BLBS H3.3 point mutants.

## Discussion

### BioID as a Tool to Study BLBS and non-BLBS Histone Point Mutants

Histones influence local chromatin states via their PTMs and the recruitment of proteins that bind them. Histone proteomes were recently probed by BioID since it allows for the detection of proximal protein associations as they occur in live cells^30,35-37^. Here we employed the BioID2 enzyme that, unlike the first-generation BirA*, lacks a DNA-binding domain^38^—a feature potentially important to this study, even though BirA* successfully identified histone associations in the aforementioned studies. The inducibly expressed H3.3 BioID2 fusion proteins represented ∼20% of the total H3.3 pool, were deposited on chromatin, and acquired PTMs (**Suppl. Fig. 1, Fig. 5**). This relative proportion of the exogenous mutant histone is theoretically comparable to that of cycling BLBS cells, where one copy out of 4 alleles is affected. However, the *H3-3A* and *H3-3B* genes are thought to have slightly different expression profiles during development^52,53^, and postmitotic cells such as neurons are particularly affected by the mutant histones^30,54^. Our model system may not be representative of post-mitotic cells, but BioID using the Flp-In T-REx system does identify disease-relevant changes in protein associations on chromatin^30,55^. Nearly two-thirds of the H3.3 prey proteins were identified regardless of the position of the biotin ligase. However, some differences were also noted, with PWWP notably being better detected (i.e., higher average spectral counts) with the biotin ligase at the H3.3 C-terminus, near the nucleosome dyad.

The PWWP deregulation was notable but likely underestimated since some proteins were barely detected, regardless of the BioID2 position. ZMYND11 seemed to be strongly affected by the point mutants, but that data is based on low spectral counts (**Fig. 4b**). Other potentially deregulated PWWP-containing proteins were not detected or failed to meet our rigorous thresholds (i.e., BFDR ≤ 1%, min. AvgSpec ≥10), namely, BRPF3, HDGFL1, HDGFL3, MBD5, PWWP2B, ZCWPW1, and ZCWPW2. Further analyses will be needed to determine how those proteins are affected by the BLBS point mutants in this study.

### Molecular Convergence on H3.3K36 and PWWP Proteins

BLBS patients display variable phenotypic severities but share core features, namely craniofacial anomalies, developmental delay, intellectual disability, and, frequently, a failure to reach developmental milestones^56^. The altered H3.3K36 methylation and PWWP associations for BLBS point mutants near glycine 34 and lysine 36 are the first common molecular feature to be shown. As with the patients, there was also variation at the molecular level with S31F, T45I causing increased K36me and PWWP binding, and G34R/V, K36E, and L48R causing a decrease.

Changes for G34R/V/W and DNMT3A/B binding were previously shown by us^30^. We now show that these mirror their impact on H3K36 methyltransferases, with the arginine residue causing the greatest impairment of catalytic activity, and valine and tryptophan having progressively milder effects (**Fig. 5d, Suppl. Fig. 8b**).

The difference between H3.3 G34V and K36E was peculiar, given the strongest overall impact of the former on PWWP proteins (**Fig. 4**). However, a closer examination of the affected proteins does reveal a similar trend, with weaker associations between the same PWWP proteins and both point mutants (**Suppl. Fig. 7**). These were likely the result of altered H3.3K36me levels since our modeling predicts a limited impact of both mutants on PWWP binding (**Fig. 4d**). Further impacts on protein binding independently of PTM are also possible. For example, while G34R/V causes steric hindrance with H3K36 methyltransferases in cis^20^, K36M preferentially interacts with the enzymes, leading to an effect in trans^19,57^.

### Mechanistic Variability in K36 Methyltransferase Deregulation

The molecular variability behind the altered H3.3K36 methylation was noteworthy (**Fig. 6**). As explained, G34R/V impair interactions with the SET domain of K36 methyltransferases. Amino acid substitutions at lysine 36 remove their target substrate, but the BLBS mutations at positions 45 and 48 stood out since both are within the H3.3 N-terminal α-helix that enters the nucleosome core^58^.

The published cryo-EM structure of NSD3 C-terminal half engaging a nucleosome was particularly insightful since it revealed extensive contacts between the N-terminal α-helix of H3 and the AWS domain^51^. The enzyme interacts with multiple nucleosomal surfaces, affecting the placement of nucleosomal DNA^51^.

Hydrophobic residues on the AWS were shown to constrain side-chain conformation on the N-terminal α-helix^59^. Select (non-BLBS) H3 and AWS amino acid substitutions altered their interaction, affecting the catalytic activity of the neighboring SET, but not the binding affinity of the enzyme fragment to its substrate^51^. It was then hypothesized that the changes to the AWS domain affected the insertion of the histone tail into the catalytic site^51^.

We were thus particularly interested in hydrophobic changes stemming from the BLBS point mutants and similarly attribute in part the impact of T45I and L48R on K36 methyltransferases to altered hydrophobic interactions with the AWS domain. The increased and decreased hydrophobicity were respectively linked to the gained H3K36me with the T45I mutant and its loss with L48R (**Fig. 5e**). This altogether meant that, unlike G34V and K36E, the T45I and L48R mutants altered H3.3K36 methylation through completely different means.

T45 and L48 alanine point mutants were also described in *S. cerevisiae*^60-62^. Curiously, the *S. cerevisiae* H3 T45A negatively impacted the lysine 36 trimethylation mark^62^, again indicating that the nature of the amino acid substitution (e.g., T45A vs T45I bulkiness) can have additional effects on K36 methyltransferases (e.g., abolish vs stimulate activity). Just as interesting, the L48A missense mutation was found to be lethal^61^.

The molecular impact of S31F remains to be established, but it increased both H3.3K36me3 and PWWP binding (**Fig. 5a, Suppl. Fig. 7**). The potential stimulation of K36 methyltransferases by that mutant will require further examination, given that H3.3S31 phosphorylation engages and stimulates the catalytic region of SETD2, ejecting the PWWP-containing ZMYND11 transcriptional regulator^63^.

Our work revealed some of the first mechanistic insights into a set of BLBS point mutants and their convergence on K36 methylation. It is, however, important to remember that our analysis identified numerous other altered protein associations (**Suppl. Figs 2-4**), and the impact of each point mutant is likely compounded. There were indeed notable changes in the binding of specific histone chaperones, not only with the G90R mutant but also with G34V, K36E, T45I, L48R, D81H, and R83C. The loss of DAXX associations with G90R is, for example, expected to change the genome-wide distribution of the mutant histone, thereby further altering its modification by histone-modifying enzymes, including K36 methyltransferases. Such a mechanism may perhaps further exacerbate the impact of the L48R missense mutation, since it also lost associations with DAXX, though to a much lesser extent (**Suppl. Fig. 4**).

Altogether, the observed changes due to BLBS point mutants lead to a model where distinct molecular effects converge on enzymes affecting H3.3K36me3 deposition in cis. Some BLBS H3.3 mutations directly affect interactions with the catalytic SET domain (e.g., G34V, K36E) while others indirectly affect SET activity through altered associations with the AWS domain (e.g., T45I, L48R). Likely, S31F functions in a yet distinct manner, and changes in DAXX binding may further compound the impacts on histone PTMs (e.g., G90R, L48R). In any case, the altered H3.3K36 methylation directly impacts the recruitment of important PWWP proteins such as DNMT3A/B.

## Acknowledgments

The work was funded by the Canadian Institutes of Health Research (CIHR RN508231 - 497056 awarded to EIC and CCLC), the Natural Sciences and Engineering Research Council of Canada (NSERC RGPIN-2024-06709 awarded to EIC), the Cancer Research Society (CRS 942402 awarded to EIC), and University of Toronto Open Scholarships awarded to SMM.

## Attributions

SMM and EIC participated in the project conception, sample preparation, data acquisition, data analysis, and manuscript preparation. SS, SV, and JFC performed the crystallization and solved the structure of the PWWP domain and ran the in vitro methyltransferase assays. CJW and ACG led the mass spectrometry experiments. AS, AM, OB, RA, GLB provided experimental assistance. JFG, NJ, CCLC, EJB, ACG, JFC, and EIC provided project and supervisory guidance.

## Data availability

Mass spectrometry data was deposited in the MassIVE repository and assigned the accession number MSV000099029. The ProteomeXchange accession is PXD068078. The PWWP – histone structure was deposited and assigned PDB ID 9Y5P.

## Supplemental Figures

**Suppl. Figure S1.**
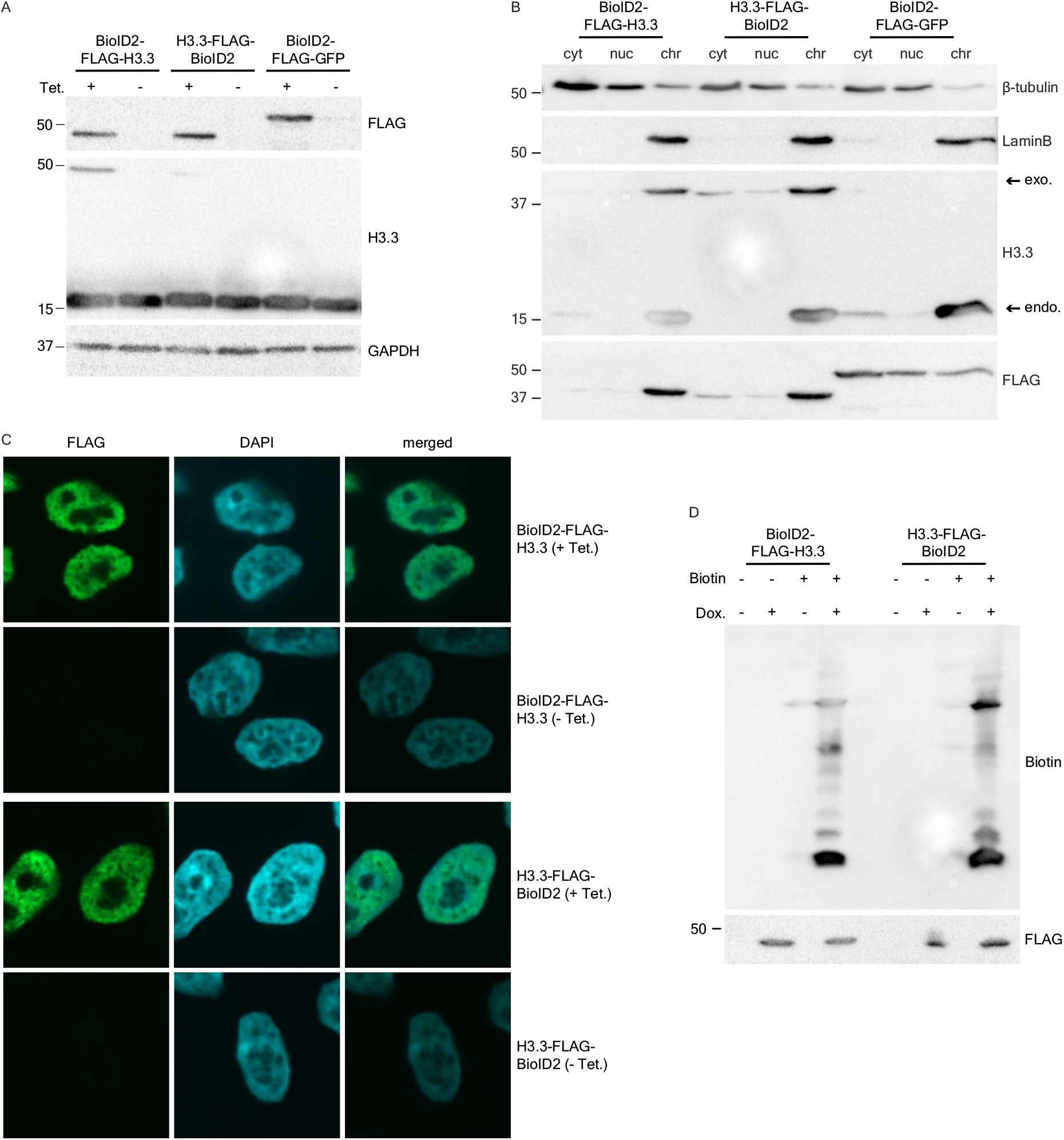
BioID controls. **(a)** Inducible expression of WT H3.3 with BioID2 fused to either of its N- or C-termini. **(b)**Subcellular fractionation showing predominant endogenous and exogenous H3.3 signals within the insoluble chromatin fraction. cyt = cytoplasmic fraction; nuc = soluble nuclear fraction; chr = insoluble chromatin fraction **(c)** Immunolabeling of exogenous (FLAG-tagged) BioID2-H3.3 constructs after a 24-hr induction period. The soluble material was extracted before labeling. **(d)** Western blot showing biotinylation of H3.3 proximal proteins upon BioID2-H3.3 induction and biotin supplementation in the media.

**Suppl. Figure S2.**
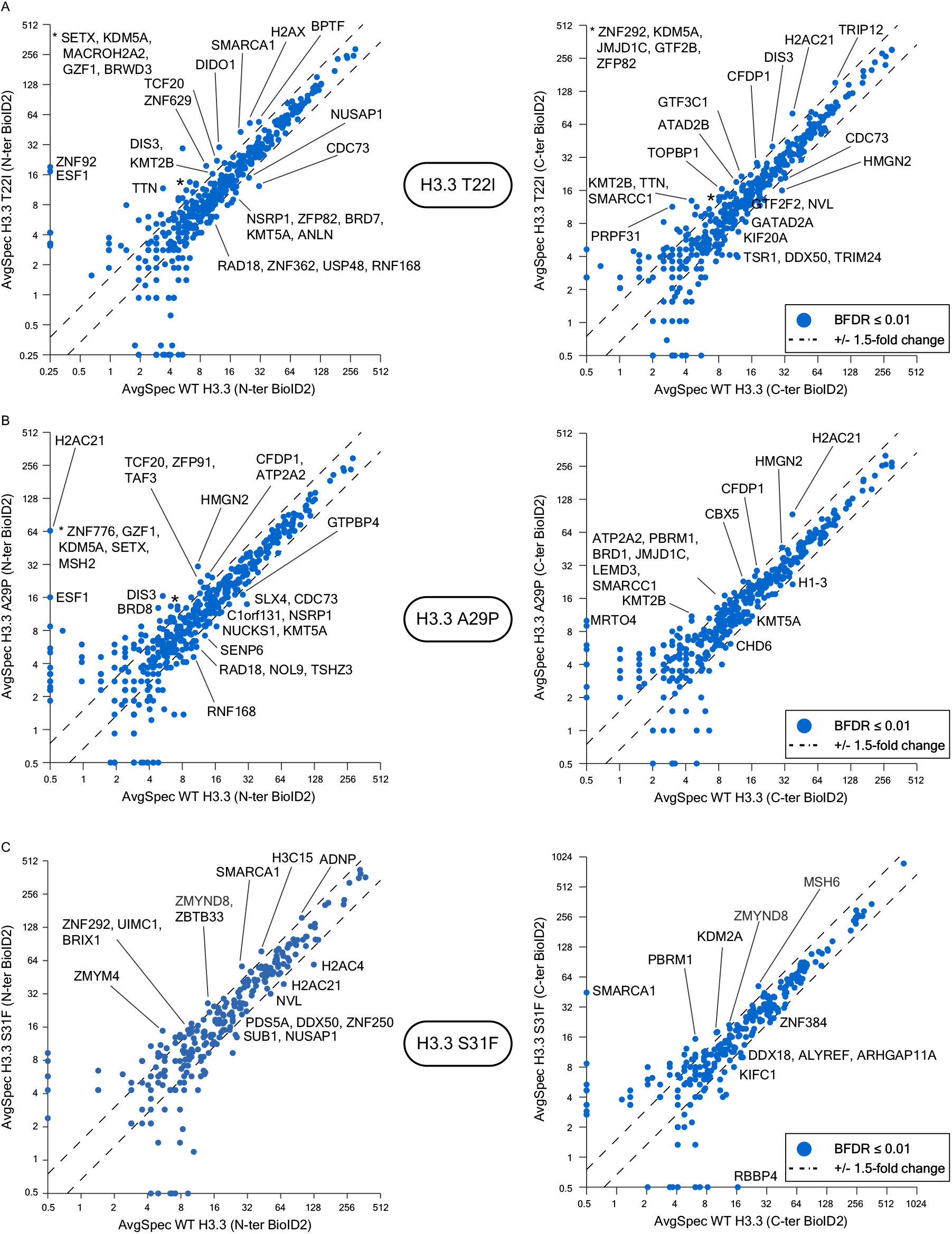
Impact of H3.3 T22I, A29P, and S31F on histone proximal associations. Scatter plots comparing average spectral counts (AvgSpec) of preys identified by **(a)** H3.3 T22I, **(b)** A29P, **(c)** S31F, and WT H3.3. Preys with ≥ 1.5-fold change and AvgSpec ≥10 on at least one condition are labelled. BioID using the N-ter BioID2 bait is shown on the left, and the one using the C-ter BioID2 bait is represented on the right.

**Suppl. Figure S3.**
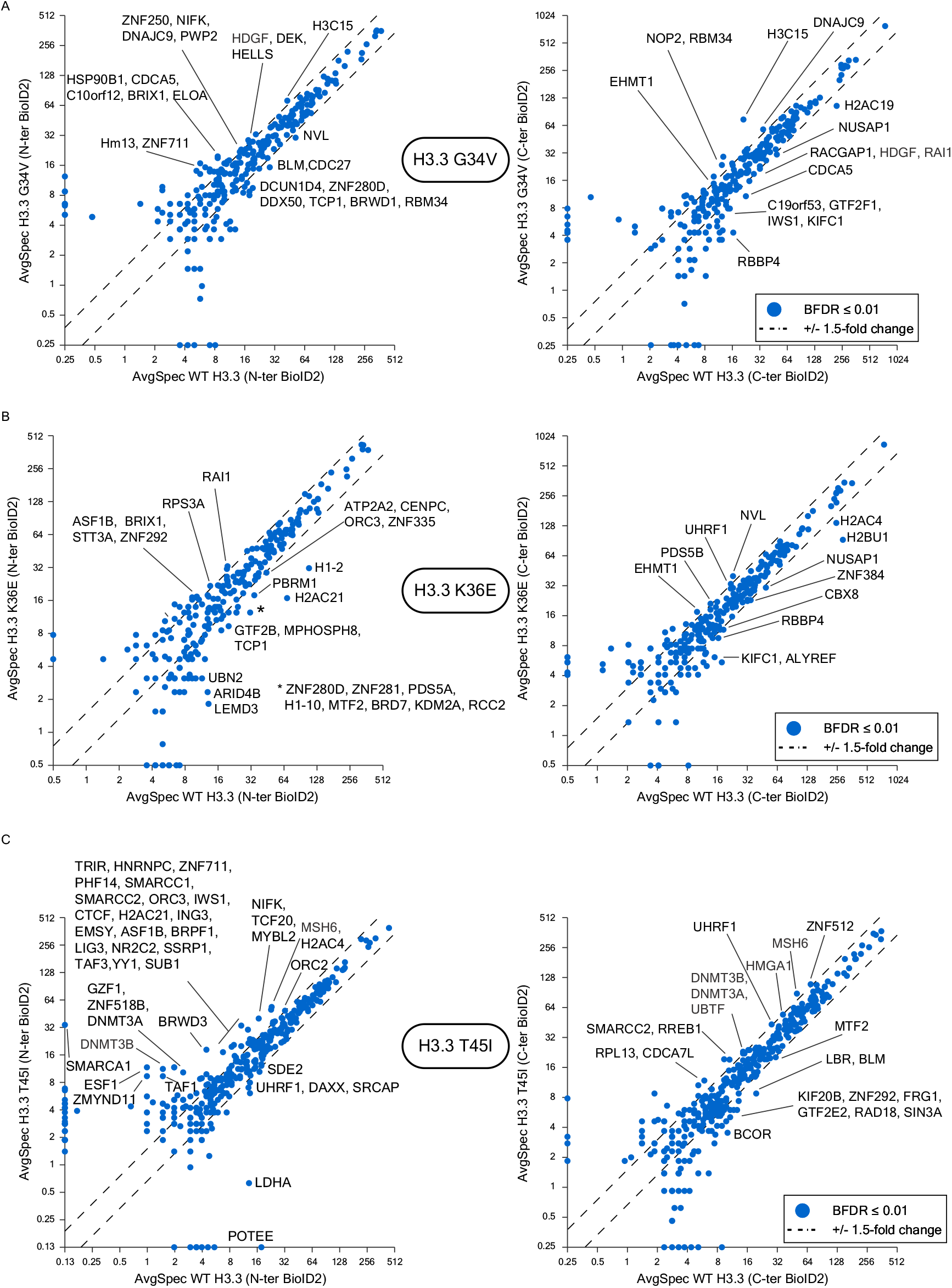
Impact of H3.3 G34V, K36E, and T45I on the histone proximal associations. Scatter plots comparing average spectral counts (AvgSpec) of preys identified by **(a)** H3.3 G34V, **(b)** K36E, **(c)** T45I, and WT H3.3. Preys with ≥ 1.5-fold change and AvgSpec ≥10 on at least one condition are labelled. BioID using the N-ter BioID2 bait is shown on the left, and the one using the C-ter BioID2 bait is represented on the right.

**Suppl. Figure S4.**
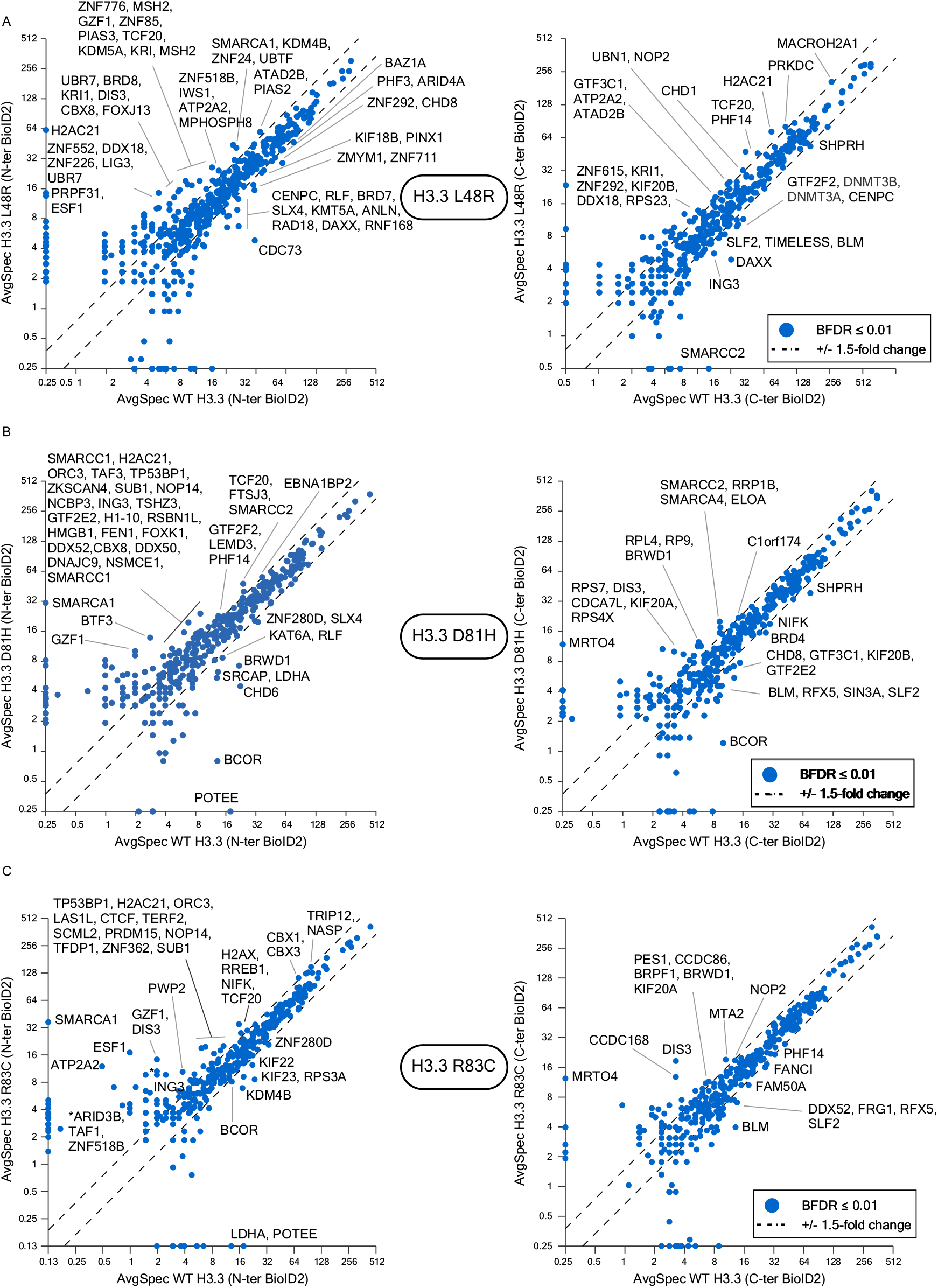
Impact of H3.3 L48R, D81H, and R83C on the histone proximal associations. Scatter plots comparing average spectral counts (AvgSpec) of preys identified by **(a)** H3.3L48R, **(b)** D81H, **(c)** R83C, and WT H3.3. Preys with ≥ 1.5-fold change and AvgSpec ≥10 on at least one condition are labelled. BioID using the N-ter BioID2 bait is shown on the left, and the one using the C-ter BioID2 bait is represented on the right.

**Suppl. Figure S5.**
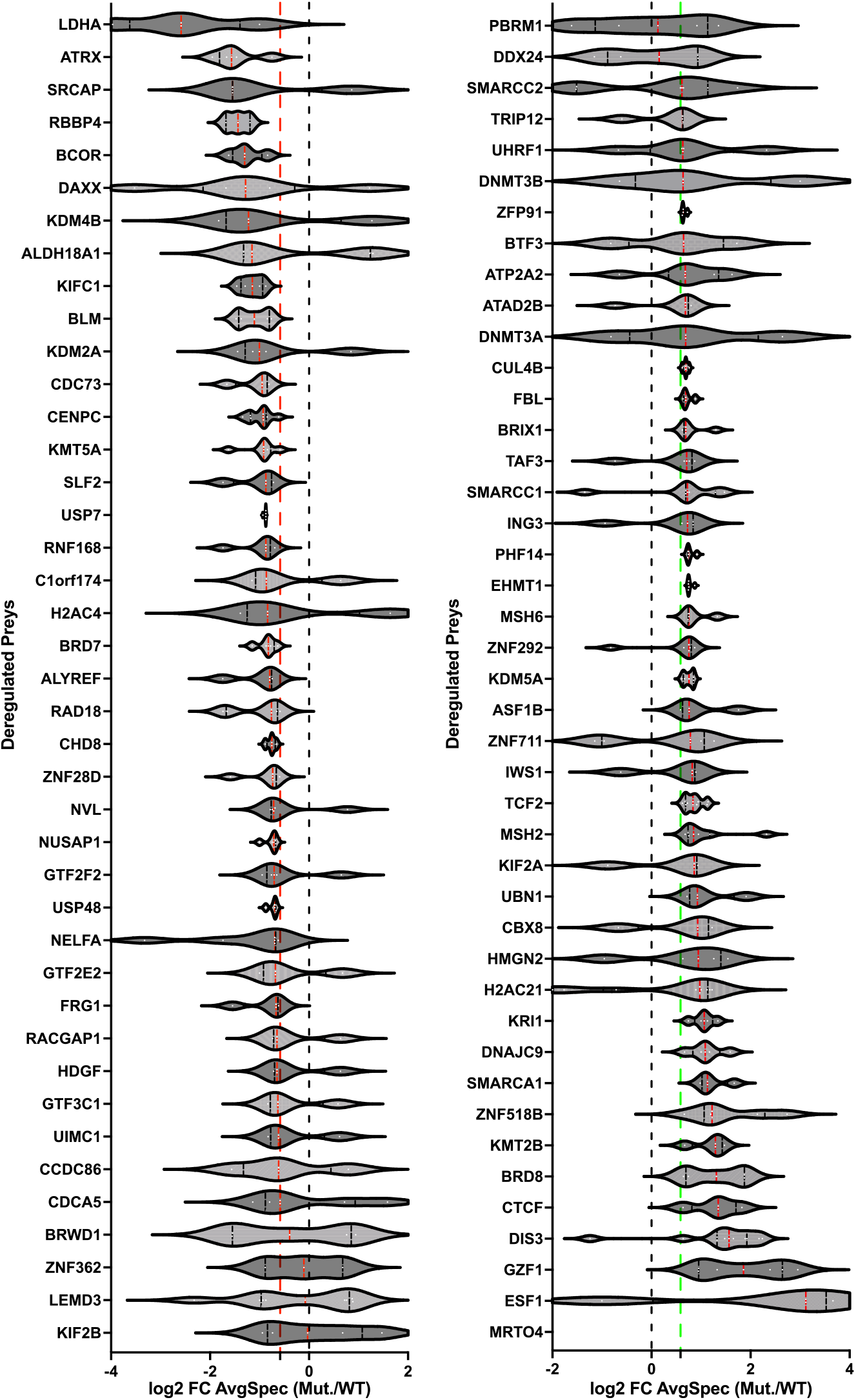
Commonly deregulated prey amongst the H3.3 mutant baits. **(a-b)** Violin plot showing the distribution of mutant/WT AvgSpec ratios for BioID prey with increased protein associations (left) and decreased ones (right). Datapoints include values from all baits (N- and C-ter tagged H3.3).

**Suppl. Figure S6.**
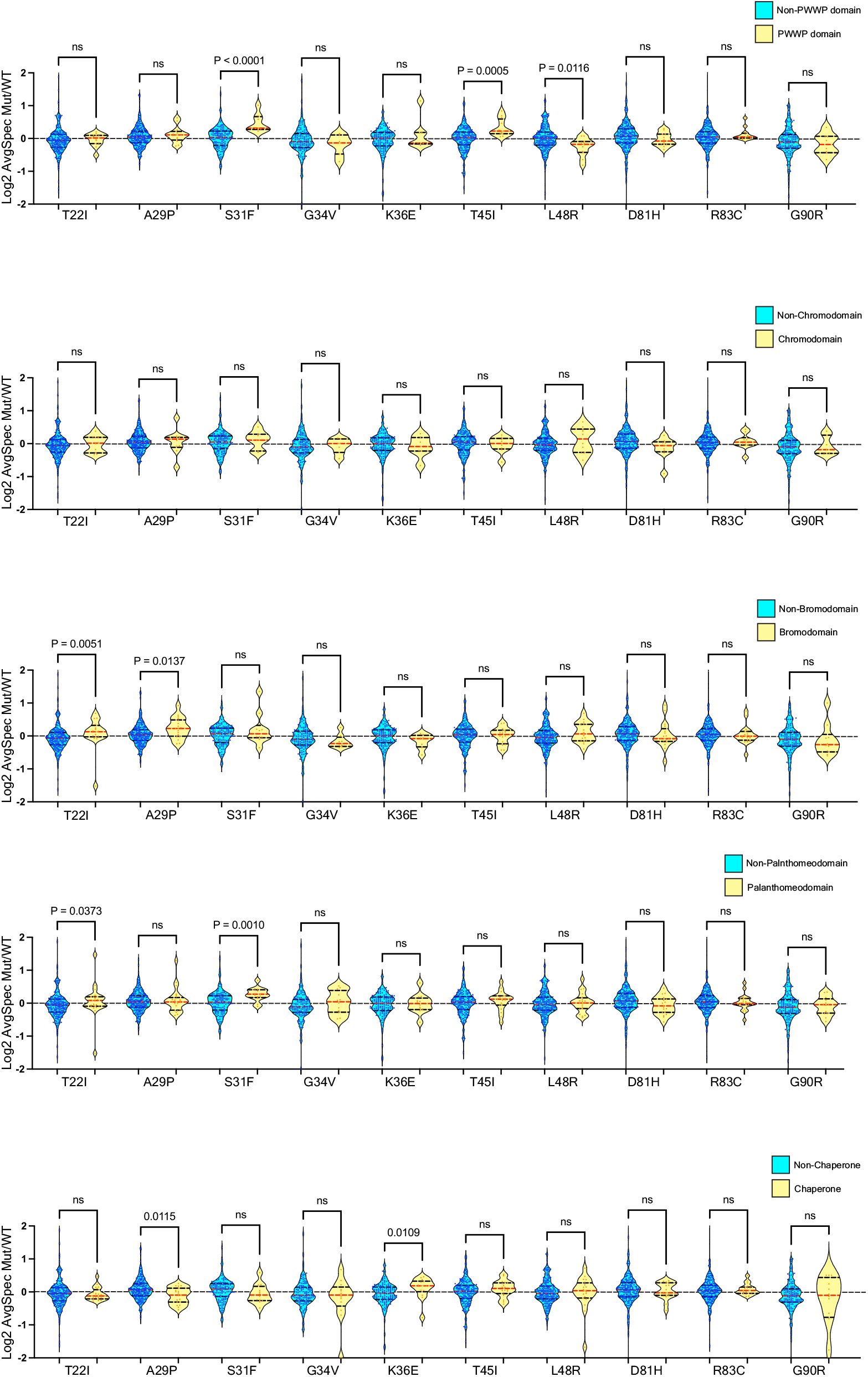
Distribution of mutant/WT AvgSpec ratios for prey with the most frequently observed histone-binding domains amongst all of the WT and mutant H3.3 BioID experiments. The last violin plot similarly shows the distribution of the mutant/WT ratios for histone chaperones. Results of a Mann-Whitney statistical analysis are shown.

**Suppl. Figure S7.**
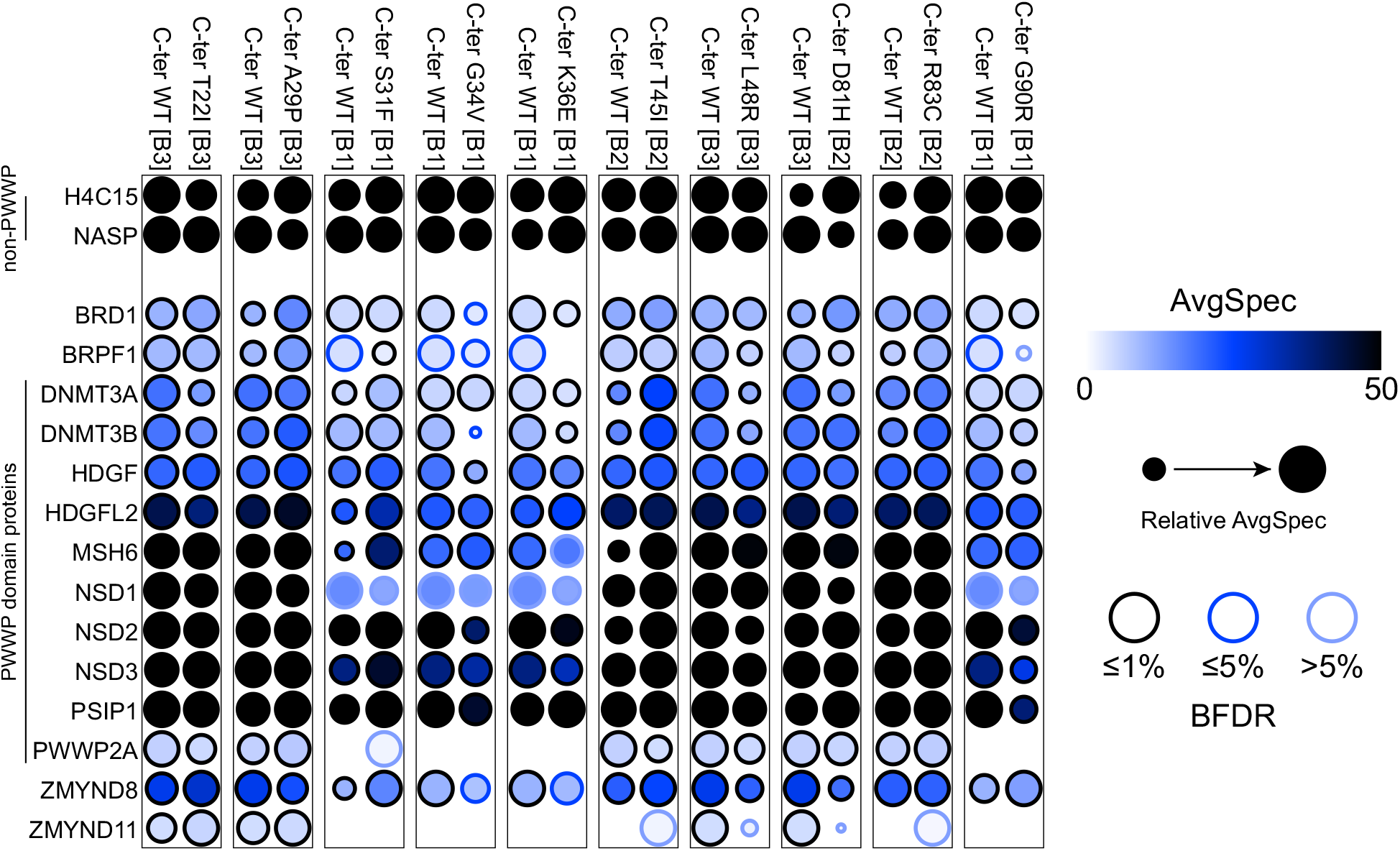
Dot plot showing prey proteins with a PWWP domain identified with the H3.3 prey.

**Suppl. Figure S8.**
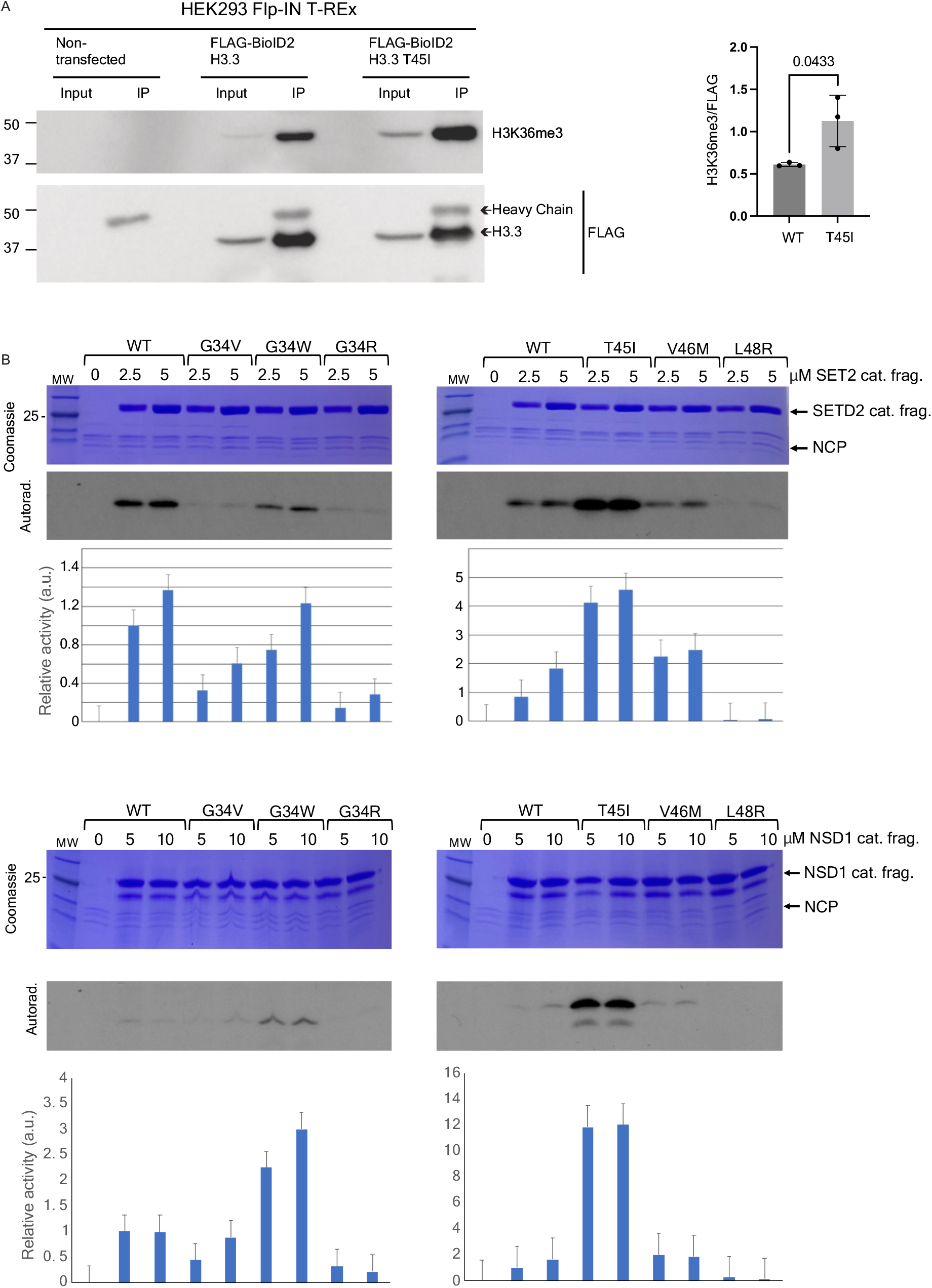
BLBS H3.3 mutants directly impact H3K36 methylation. **(a)** Western blot analysis of FLAG-immunoprecipitated N-terminally tagged FLAG-BioID2 H3.3 (WT and H3.3T45I). Inputs represented 5% of the extracts. IP = immunoprecipitation. An unpaired t-test statistical analysis compared the H3K36me3/FLAG signal of H3.3T45I with WT H3.3 values from prey proteins with and without the protein domains (n=3). **(b)** In vitro methyltransferase assays. (n=3-4 for SETD2, and n=2 for NSD1)

**Suppl. Table S1.** Protein domains recurrently found within H3.3 prey proteins. Data was obtained using the ProHits-viz^66^ domain enrichment analysis tool and the InterPro database^67^.

| Domain | Interpro identifier | # of preys |
| --- | --- | --- |
| ADD domain | IPR025766 | 3 |
| Ankyrin repeat | IPR002110 | 9 |
| ARID DNA-binding domain | IPR001606 | 7 |
| AWS domain | IPR006560 | 5 |
| Basic-leucine zipper domain | IPR004827 | 4 |
| BEN domain | IPR018379 | 6 |
| BRCT domain | IPR001357 | 11 |
| Bromo adjacent homology (BAH) domain | IPR001025 | 7 |
| Bromodomain | IPR001487 | 29 |
| Chromo/chromo shadow domain | IPR000953 | 16 |
| DDT domain | IPR018501 | 5 |
| Extended PHD (ePHD) domain | IPR034732 | 12 |
| G-patch domain | IPR000467 | 18 |
| Helicase superfamily 1/2, ATP-binding domain | IPR014001 | 58 |
| Helicase, C-terminal domain-like | IPR001650 | 59 |
| High mobility group box domain | IPR009071 | 17 |
| Homeodomain | IPR001356 | 10 |
| JmjC domain | IPR003347 | 2 |
| Krueppel-associated box | IPR001909 | 48 |
| Linker histone H1/H5, domain H15 | IPR005818 | 6 |
| MCM domain | IPR001208 | 6 |
| Methyl-CpG DNA binding | IPR001739 | 7 |
| PDZ domain | IPR001478 | 10 |
| Post-SET domain | IPR003616 | 11 |
| Pre-SET domain | IPR007728 | 3 |
| Protein kinase domain | IPR000719 | 20 |
| PWWP domain | IPR000313 | 16 |
| RNA helicase, DEAD-box type, Q motif | IPR014014 | 21 |
| SANT domain, SANT/Myb domain | IPR017884, IPR001005 | 22 |
| SAP domain | IPR003034 | 12 |
| SET domain | IPR001214 | 20 |
| Tudor domain | IPR002999 | 2 |
| WD40 repeat | IPR001680 | 31 |
| WW domain | IPR001202 | 11 |
| Zinc finger C2H2-type | IPR013087 | 128 |
| Zinc finger, CCCH-type | IPR000571 | 16 |
| Zinc finger, CXXC-type | IPR002857 | 4 |
| Zinc finger, PHD-finger | IPR019787 | 37 |

## Online Methods

### Expression vectors

Mouse H3.3 cDNA encoding a protein identical to the human H3.3 was PCR amplified from pETa-H3.3 using primers flanked by attB sequences. Reverse primers generated amplicons with or without a stop codon (referred here as closed or open constructs, respectively). The amplified DNA was introduced into pDONR223 through a Gateway BP reaction following the manufacturer’s protocol (ThermoFisher Scientific). The resulting pDONR223-H3.3-closed and pDONR223-H3.3-open plasmids were subject to site-directed mutagenesis per a modified QuickChange protocol (Agilent) to generate pDONR223-H3.3 with BLBS mutations. An LR reaction subsequently introduced the different H3.3 constructs into pcDNA5 destination plasmids generated by Dr. Anne-Claude Gingras (Lunenfeld-Tanenbaum Research Institute) to obtain pcDNA5-BioID2-FLAG (N-ter tagged H3.3) and pcDNA5-FLAG-BioID2 (C-ter tagged H3.3). All plasmids were verified by Sanger sequencing (TCAG Facility, SickKids Research Institute).

Site-directed mutagenesis was performed as described above to generate pETa-H3.3 vectors with the G34R/V/W, T45I, V46M, and L48R mutations. The pFastBac-NSD1-FL and pFastBac-SETD2-FL vectors were previously generated by Hwa Young (Haley) Yun in the Campos Lab and were used to PCR amplify the AWS, SET, and postSET-encoding regions and introduced into a pGEX vector (MilliporeSigma). DNMT3B (3QKJ) was a gift from Dr. Cheryl Arrowsmith, PMCC and Structural Genomics Consortium (SGC) at the University of Toronto (Addgene plasmid # 32044).

### Stable cell lines and induced expression

HEK293 Flp-In T-REx cells (ThermoFisher Scientific) were cultured in DMEM (Wisent Inc.) supplemented with 10% FBS (Wisent Inc.), 1x penicillin/streptomycin (Wisent Inc.), 75 µg/ml zeocin (Invitrogen), 5 µg/ml blasticidin (Invivogen), and kept subconfluent through regular passaging. 1×10^6^ cells were seeded in media without zeocin or blasticidin and co-transfected using Lipofectamine 3000 (ThermoFisher Scientific) with the Flp recombinase-encoding pOG44 (ThermoFisher Scientific) and one of the following: pCDNA5 H3.3-FLAG-BioID2 (WT or mutant), pCDNA5 BioID2-FLAG-H3.3 (WT or mutant), or pCDNA5 GFP-BioID2 in a 9:1 ratio. A negative control was only transfected with pOG44. Fresh media without selection drugs was supplied 24 hours following transfection and replaced with media supplemented with 200 µg/ml hygromycin (Wisent Inc.) and 5 µg/ml blasticidin (Invivogen) 48 hrs post-transfection. Media was changed periodically, and stable clones selected through continued hygromycin exposure for at least one week or until the negative control showed no live cells. The hygromycin concentration was reduced to 100 µg/ml once this was achieved. Cells were induced with 1ug/ml doxycycline for 24 hours.

### Antibodies

Antibodies raised against the following were used: Biotin (Cell Signaling Technologies 5597), DAXX (Santa Cruz sc-7152), mouse anti-FLAG (MilliporeSigma F1804), rabbit anti-DDDDK (Abcam ab205606), H3 (Abcam ab1791), H3.3 (Abcam ab176840), mouse anti-H3K36me3 (Active Motif 61021), rabbit anti-H3K36me3 (Abcam ab9050), LMNB1 (Proteintech 66095), and β-tubulin (DSHB E7).

### Western blotting

Whole cell extracts were obtained by resuspending cells in RIPA lysis buffer (50 mM tris pH 7.4, 150 mM NaCl, 1% NP-40, 0.1% SDS, 1 mM EDTA, and 0.5% sodium deoxycholate) with fresh protease inhibitors, followed by gentle sonication thrice 10 seconds. Whole cell extracts were incubated at 4 °C overnight with 0.25 units/µL MNase in the presence of 5 mM CaCl_2_ and additional PMSF. The concentration of the whole cell extracts was determined using a Bradford reagent (Bio-Rad). 25 to 50 µg of the cell extracts were briefly boiled in 1X Laemmli buffer, loaded onto a 15% polyacrylamide gel, resolved by electrophoresis, and transferred onto activated PVDF membranes (Amersham Hybond). Membranes were blocked in 5% milk for 1 hour before rinsing twice in TBST (1 mM Tween 20, 20 mM tris pH 7.5, 150 mM NaCl) and overnight incubation in primary antibody. Membranes were washed 3x 5 minutes in TBST before incubating with HRP-coupled secondary antibodies (Rockland Immunochemicals) for 1 hour at room temperature, followed by 3x 5 minutes washes and band visualization with a luminol solution (100 mM tris pH 8.8, 1.25 mM luminol, 0.2 mM coumaric acid, activated with 5 µl 10% H_2_O_2_ per 1 ml) using Bio-Rad ChemiDoc MP. Western signals were quantified using Image Lab (Bio-Rad), and statistical analyses were done using Prism (GraphPad).

### Immunoprecipitation

Cells were resuspended in 100-200 µl IP lysis buffer (50 mM tris pH 7.6, 100 mM KCl, 2 mM EDTA, 0.1% NP40, 10% glycerol, supplemented with fresh proteinase inhibitors), flash frozen on dry ice for 15 minutes, and thawed at room temperature. Whole cell extracts were treated with 0.25 units/µL MNase in the presence of 5 mM CaCl_2_ for a minimum of 3 hours at 4 °C. For each mg of protein, 5 µl ProteinG magnetic beads (GE Healthcare) were washed 3x with 500 µl IP lysis buffer at 4 °C and resuspended in 100 µl IP + 2 µg primary antibody and incubated with gentle rotation for a minimum of 3 hours at 4 °C. The antibody-coupled beads were then incubated with the cell extracts. Immunoprecipitations were carried out overnight at 4 °C, after which beads were washed 3x with IP lysis buffer without NP40. Beads were resuspended in 1x Laemmli buffer, boiled, and analyzed by western blotting.

### Subcellular fractionation

Cells were gently resuspended in 2 pellet volumes of cytosolic extraction buffer (20 mM tris pH 7.6, 10 mM NaCl, 1 mM EDTA, 0.5% NP-40, 0.5 mM DTT, and fresh protease inhibitors) and immediately spun for 5 minutes at 500 x *g* at 4 °C to pellet the nuclei and organelles. The supernatant (cytoplasmic fraction) was stored at −80°C after adjusting the salt concentration to 100 mM and supplementing with 20% glycerol (final concentration). Pellets were resuspended in 3 pellets volume of nuclear extraction buffer (20 mM tris pH 7.6, 300 mM NaCl, 1 mM EDTA, 20% glycerol, 0.5 mM DTT, and fresh protease inhibitors) and centrifuged at 30,000 x *g* for 15 minutes at 4 °C. The supernatant (soluble nuclear extract) was stored at −80 °C after adjusting the salt concentration to 100 mM NaCl by diluting samples with BC0 (20 mM tris pH 7.6, 0.2 mM EDTA, 20% glycerol, 0.5% DTT, and fresh protease inhibitors). Pellets, containing the insoluble nuclear fraction, were resuspended in BC100 (BC0 with 100 mM NaCl) and sonicated ∼3x 15 seconds (until there were no visible clumps). Chromatin was digested by incubating samples with 0.25 units/µl MNase in the presence of 5 mM CaCl_2_ with fresh PMSF overnight at 4 °C. 25 µg of each fraction were then resolved on a 15% PAGE.

### Histone Acid extraction

A protocol based on Shechter D. et al^68^ was used. Briefly, pellets containing ∼20×10^6^ cells were resuspended in 1 ml of ice-cold hypotonic lysis buffer (10 mM Tris-Cl pH 8.0, 1 mM KCl, 1.5 mM MgCl_2_) supplemented with proteinase inhibitors. Extraction was performed at 4 °C under gentle rotation for 1 hour. Nuclei were pelleted by centrifugation at 10,000 x *g* for 10 minutes at 4°C. Supernatant was discarded and nuclei were resuspended in 0.8 ml 0.4N H_2_SO_4_ and incubated for at least 1 hour at 4 °C with gentle rotation. Nuclear debris was removed by centrifugation at 16,000 x *g* for 10 minutes at 4 °C. The supernatant containing histones and other acid-soluble nuclear proteins was transferred to a fresh tube and precipitated by the addition of 0.264 ml 6.1N trichloroacetic acid (TCA), final concentration of 1.5N TCA. Samples were precipitated for at least 1 hour with gentle rotation. The precipitated proteins were pelleted by centrifugation at 20,000 x *g* for 10 minutes at 4 °C, and the supernatant was discarded. Precipitated proteins were washed with 1 ml 100% acetone thrice, with centrifugation in between washes at 20,000 x *g* for 10 minutes at 4 °C. Washed proteins were air-dried for 20-30 minutes at room temperature, followed by resuspension in BC100. Samples were gently rotated at room temperature for 1 hour. Any undissolved material was removed by centrifugation at 20,000 x *g* for 10 minutes at 4 °C. The acid-extracted proteins were stored at −80 °C.

### Immunofluorescence

Coverslips were coated with 1:10 diluted poly-L-lysine (MilliporeSigma) for 10 minutes at room temperature with occasional swirling, after which coverslips were rinsed with water and air-dried. HEK293 Flp-In T-REx cells were seeded at 0.25-0.5×10^6^ cells per well. After reaching ∼60% confluency, they were induced with 1 µg/ml doxycycline for 24 hours. Media was aspirated, and cells fixed with a filtered 2% paraformaldehyde solution containing 0.2% triton X-100 (MilliporeSigma), pH 8.2 in ddH_2_O (Electron Microscopy Sciences) at room temperature for 20 minutes. Cells were then rinsed 3x 5 minutes with PBS with slow rocking at room temperature, and then permeabilized with filtered 0.5% TERGITOL solution (MilliporeSigma) in PBS, for 10 minutes at room temperature. Coverslips were again rinsed 3x 5 minutes with PBS and antigens blocked using a blocking buffer composed of 3% bovine serum albumin (Multicell) and 1% normal goat serum (Rockland Immunochemicals) diluted in PBS for 1 hour at room temperature. Coverslips are then moved to a humidity chamber and incubated with primary antibody diluted in blocking buffer for 1 to 2 hours at room temperature. Coverslips were rinsed 3x 5 minutes each with PBS. Alexa Fluor 594-conjugated anti-rabbit (Jackson Immunochemicals) and Alexa Fluor 647-conjugated anti-mouse secondaries (BioLegend) were diluted in blocking buffer to incubate coverslips in the humidity chamber for 30 minutes at room temperature. Coverslips were again rinsed 3x 5 minutes each with PBS and incubated with DAPI diluted in PBS with a final concentration of 1 µg/ml for 10 minutes at room temperature in the humidity chamber. Coverslips were rinsed a final time for 5 minutes with PBS before being mounted on glass slides using ProLong Gold antifade reagent (Invitrogen) and allowed to cure overnight at 4°C.

### BioID Sample Preparation

All BioID samples were prepared through independent duplicates. Each cell line was seeded in 15 cm plates, grown under normal conditions until 60-70% confluency was reached. Cells were induced for 24 hours with 1 µg/ml doxycycline, and the media was supplemented with 50 µM biotin (final concentration) during the last 8 hours of induction. Cells were then washed with cold PBS, scraped, spun down at 500 x *g* at 4 °C for 5 minutes, and flash frozen after removal of the supernatant. The induction and biotinylation were confirmed from a small sample aliquot by western blotting.

Frozen cell pellets were resuspended in ice-cold modified RIPA lysis buffer (50 mM tris pH 7.4, 150 mM NaCl, 1 mM EGTA, 0.5 mM EDTA, 1 mM MgCl2, 1% NP40, 0.1% SDS, 0.4% sodium deoxycholate, 1 mM PMSF (fresh), protease inhibitors (fresh)) at a 1:4 pellet weight:volume ratio. Cells were sonicated for 15 sec (5 sec on, 3 sec off for three cycles) at 30% amplitude on a sonicator with a 1/8” microtip while kept on ice. 250 U of TurboNuclease and 10 µg of RNase were then added, and samples were rotated (end-over-end) at 4°C for 15 min. SDS concentration was increased to 0.4% (by the addition of 10% SDS), and samples were rotated at 4°C for 5 min. Samples were centrifuged at 20,817 x *g* for 20 min at 4°C, and the supernatant was transferred to a 2 ml centrifuge tube.

A master mix of streptavidin beads using 35 µl of slurry (20 µl bed volume) for each sample, plus 10% to account for any losses, was prepared. Streptavidin agarose beads were washed 3 times with 1 ml lysis buffer (minus proteinase inhibitors, PMSF, and deoxycholate). Following the last wash, beads were resuspended as a 50% slurry. 40 µl of the 50% slurry of streptavidin beads was added to the clarified supernatant and rotated using gentle end-over-end rotation for 3 hours at 4°C. Beads were pelleted by centrifugation at 400 x *g* for 1 min at 4°C. Supernatant was discarded. Beads were transferred in 1 ml fresh RIPA-wash buffer (50 mM Tris-HCl, pH 7.4, 150 mM NaCl, 1 mM EDTA, 1% NP40, 0.1% SDS, 0.4% sodium deoxycholate) to a new microcentrifuge tube. Beads were washed once with SDS-Wash buffer (25 mM Tris-HCl, pH 7.4, 2% SDS), twice with RIPA-wash buffer, once with TNNE buffer (25 mM Tris-HCl, pH 7.4, 150 mM NaCl, 0.1% NP40, 1 mM EDTA), and three times with 50 mM ammonium bicarbonate pH 8.0 (ABC) buffer. For each wash step, mixing was done by inversion, centrifugation at 400 x *g* for 1 min, and the supernatant was discarded by aspiration.

### Trypsin digestion

Residual ABC buffer was removed by pipette. Beads were resuspended in 70 µl of 50 mM ABC buffer with 1 µg trypsin, also dissolved in 50 mM ABC buffer. Samples were incubated at 37°C overnight with agitation. An additional 0.5 µg of trypsin was added, and the samples were incubated for a further 3 hours. Beads were centrifuged at 400 x *g* for 2 min, and the supernatant was collected in a new 1.5 ml tube. Beads were washed twice with 150 µl mass spectrometry grade H20 (pelleting beads in between), and the wash supernatant was added to the peptide supernatant. The supernatant was centrifuged at 16,100 x *g* for 10 min, and most of it was transferred to a new tube (leaving ∼30 µl residual so as not to transfer beads). The pooled supernatant was lyophilized using vacuum centrifugation without heat. Dried peptides were stored at −80°C until ready for mass spectrometry analysis.

### Mass spectrometry analysis

For data-dependent acquisition (DDA) LC-MS/MS, affinity-purified and digested peptides were analyzed using a nano-HPLC (High-performance liquid chromatography) coupled to MS. For both anti-FLAG AP-MS and BioID, one-quarter of the sample was used. Nano-spray emitters were generated from fused silica capillary tubing, with 100 µm internal diameter, 365 µm outer diameter, and 5-8 µm tip opening, using a laser puller (Sutter Instrument Co., model P-2000, with parameters set as heat: 280, FIL = 0, VEL = 18, DEL = 2000). Nano-spray emitters were packed with C18 reversed-phase material (Reprosil-Pur 120 C18-AQ, 3 µm) resuspended in methanol using a pressure injection cell. Samples in 5% formic acid were directly loaded at 800 nl/min for 20 min onto a 100 µm × 15 cm nano-spray emitter. Peptides were eluted from the column with an acetonitrile gradient generated by an Eksigent ekspert™ nanoLC 425, and analyzed on a TripleTOF™ 6600 instrument (AB SCIEX, Concord, Ontario, Canada). The gradient was delivered at 400 nl/min from 2% acetonitrile with 0.1% formic acid to 35% acetonitrile with 0.1% formic acid using a linear gradient of 90 min. This was followed by a 15 min wash with 80% acetonitrile with 0.1% formic acid, and equilibration for another 15 min to 2% acetonitrile with 0.1% formic acid. The total DDA protocol is 120 min. The first DDA scan had an accumulation time of 250 ms within a mass range of 400–1800Da. This was followed by 10 MS/MS scans of the top 10 peptides identified in the first DDA scan, with an accumulation time of 100 ms for each MS/MS scan. Each candidate ion was required to have a charge state from 2–5 and a minimum threshold of 300 counts per second, isolated using a window of 50mDa. Previously analyzed candidate ions were dynamically excluded for 7 seconds.

### DDA Search

Mass spectrometry data generated were stored, searched and analyzed using ProHits^69^ laboratory information management system (LIMS) platform. Within ProHits^69^, WIFF files were converted to an MGF format using the WIFF2MGF converter and to an mzML format using ProteoWizard (V3.0.10702) and the AB SCIEX MS Data Converter (V1.3 beta). The data was then searched using Mascot^70^ (V2.3.02) and Comet^71^ (V2016.01 rev.2). The spectra were searched with the human and adenovirus sequences in the RefSeq database (version 57, January 30th, 2013) acquired from NCBI, supplemented with “common contaminants” from the Max Planck Institute (http://maxquant.org/contaminants.zip) and the Global Proteome Machine(GPM; ftp://ftp.thegpm.org/fasta/cRAP/crap.fasta), forward and reverse sequences (labeled “gi|9999” or “DECOY”), sequence tags (BirA, GST26, mCherry and GFP) and streptavidin, for a total of 72,481 entries. Database parameters were set to search for tryptic cleavages, allowing up to 2 missed cleavage sites per peptide with a mass tolerance of 35 ppm for precursors with charges of 2+ to 4+ and a tolerance of 0.15 amu for fragment ions. Variable modifications were selected for deamidated asparagine and glutamine and oxidized methionine. Results from each search engine were analyzed through TPP (the Trans-Proteomic Pipeline, v.4.7 POLAR VORTEX rev 1) via the iProphet^72^ pipeline.

### SAINTexpress Analysis

The SAINT analysis tool is used to identify high-confidence protein interactors versus control samples, in this case BioID2-FLAG-GFP. Prior to applying SAINT (v3.6.3), proteins were filtered based on iProphet^72^ ≥0.95 and unique peptides ≥2. iProphet^72^ cut-off was to ensure a high probability of MS2 fragmentation pattern being correctly assigned to both peptide and protein. Unique peptides cut-off was to ensure that the observed protein was the one in question. Proteins with a BFDR (Bayesian False Discovery) ≤0.01 were considered high-confidence protein interactors.

### Data analysis

Following SAINTexpress^73^ analysis, the data was run through ProHits-Viz^66^ with the default settings except the lower BFDR being adjusted to 1%. Data processed through ProHits-Viz^66^ was then used to perform subsequent analyses, from which prey detected with an average spectral count of <10 were removed (except for the condition-condition scatter plots, where at least one sample needed to meet the cutoff). Subsequently, the fold change was calculated using the ratio of the mutant bait to the WT bait. This also mitigated batch-specific effects. A fold change of ± 1.5 was chosen to indicate deregulated prey.

Protein domains found in the dataset were identified using the ProHits-Viz^66^ domain enrichment feature and the InterPro database^67^. Violin plots and Mann-Whitney statistical analyses were generated using Prism version 10.4.2 (GraphPad).

### Recombinant Proteins

Rosetta cells were transformed with His-tagged *Xenopus laevis* H2A, H2B, and H4 constructs and the mammalian H3.3 plasmids and grown in 2xYT media supplemented with kanamycin and chloramphenicol at 37°C to an OD of 0.5. Protein expression was induced by adding isopropyl-thiogalactopyranoside (IPTG, 0.4 mM) at 30°C and left overnight.

Bacteria co-expressing the histones were harvested and resuspended in octamer buffer (20 mM Tris pH8, 2M NaCl, 0.5 mM TCEP) and lysed by sonication. The lysate was centrifuged at 16,000 rpm for 30 minutes at 4°C. The supernatant was added to 10 ml of slurry talon beads, supplemented with 30 mM Imidazole (final) for 1 hour. Beads were collected and washed twice with octamer buffer supplemented with 30 mM Imidazole (final). Beads were transferred to a column and eluted with an imidazole gradient from 30 mM to 500 mM (with 40 mM increases) in octamer buffer over 12 fractions total, 2 ml each fraction, at 4°C. Fractions were loaded on 15% gel, and those that contained an equivalent amount of 4 histones were pooled and concentrated to 500 µl using an Amicon 10 MWCO. 10 µl of 1U/µl thrombin was added to digest overnight at 4°C. Octamer was further purified by gel filtration chromatography (S200 analytical) in gel filtration buffer (2M NaCl, 20 mM Tris pH 8, 0.5 mM TCEP).

Dual GST and HIS-tagged NSD1 and SETD2 pGEX-4T constructs were assembled and verified using DNA sequencing. Rosetta cells were transformed with the vectors and grown in Luria-Bertani broth media supplemented with 100 µM ZnCl_2_, ampicillin, and chloramphenicol at 37°C to an OD of 0.5. Protein expression was induced by adding isopropyl-thiogalactopyranoside (IPTG, 0.1 mM) at 37°C and left overnight.

Bacteria expressing NSD1 and SETD2 were harvested and resuspended in PBS supplemented with 500 mM NaCl, 100 µM ZnCl2, and 10 µl of SAM (10 mg/ml 100 µl aliquot) on ice, stirring for 15 minutes. Cells were then lysed by sonication and centrifuged at 16,000 RPM for 30 minutes at 4°C. The supernatant was filtered and applied onto 10 mL GST slurry beads prewashed with H2O and PBS, and left to rotate for 1 hour. Beads were collected and washed with 100 ml of PBS and resuspended in 25 ml of modified gel filtration buffer (20 mM Tris, pH 8.0, 150 mM NaCl, 5 mM βME) and TEV protease to cleave overnight at 4°C, rotating. The supernatant was collected and applied onto 5 mL slurry Talon beads. A 30 min batch binding was performed, after which eluates were concentrated and further purified by gel filtration chromatography (S75 prep) in modified gel filtration buffer.

A DNMT3B PWWP-expressing bacterial culture was incubated at 37°C with shaking at 250 rpm until an optical density at 600 nm (OD600) of approximately 1.5 was reached. At this point, the temperature was decreased to 18°C, and protein expression was induced by adding 1 mM isopropyl β-D-1-thiogalactopyranoside (IPTG) from a 1 M stock solution. Cultures were then incubated overnight at 18°C. Cells were harvested by centrifugation at 4000 rpm for 30 minutes at 4°C. Cell pellets were resuspended in 25 mL Buffer A (50 mM HEPES pH 7.4, 500 mM NaCl, 2 mM βME, 5% glycerol, 0.1% CHAPS) and stored at −80°C.

Frozen bacterial pellets corresponding to 1 liter of culture were thawed and resuspended to a final volume of 100 mL with Buffer A. Cells were lysed by sonication on ice using three cycles of 1 minute each in a stainless-steel cup. The lysate was clarified by centrifugation at 16,000 rpm for 30 minutes at 4°C. The supernatant was filtered through a 0.45 µm filter and applied onto a Ni-NTA resin column (5 mL resin) pre-equilibrated with 20 column volumes (CV) of Buffer A. The column was washed with 20 CV of Buffer A, and the bound protein was eluted with 100 mL Buffer B (50 mM HEPES pH 7.4, 500 mM NaCl, 500 mM imidazole, 2 mM βME, 5% glycerol), collecting the first 30 mL for subsequent steps. To remove the affinity tag, two vials of TEV protease were added, and the cleavage reaction was carried out overnight at 4°C in 2 L of Buffer C (50 mM HEPES pH 7.4, 250 mM NaCl, 2 mM βME). Following overnight cleavage, the sample was centrifuged at 3000 rpm for 5 minutes at 4°C to remove any precipitates. The clarified sample was transferred to a new tube and applied to a Heparin affinity column pre-equilibrated with Buffer D (50 mM HEPES pH 7.4, 2 mM βME) at a flow rate of 1 mL/min. The column was washed with 2 CV of Buffer D and eluted with a linear gradient from 0 to 100% Buffer E (0 mM to 1 M NaCl in 50 mM HEPES pH 7.4 2 mM βME). Fractions containing DNMT3A/B were pooled, concentrated to 2 mL, and further purified by size-exclusion chromatography using a Superdex 75 pre-equilibrated with 20 mM HEPES pH 7.0, 150 mM NaCl, and 5 mM βMe. Fractions 19 to 22 from the S75prep column were collected, concentrated further using a 3.5 kDa MWCO Centricon, and centrifuged at maximum speed for 5 minutes at 4°C. PWWP concentration was quantified by UV and stored at −80°C.

### Nucleosome assembly

Nucleosomes were assembled using the salt gradient dialysis method^74^. Nanodrop was used to quantify 601 DNA and histone octamer concentrations. Then, eight dilutions of nucleosome were prepared, with constant DNA concentration but increasing octamer concentration. Each dilution was supplemented with 2M NaCl. The reactions were assembled, on ice, in this order: H_2_O, NaCl, DNA, and octamer. Dilution buffer (10 mM Tris pH8) was added every 30 minutes at room temperature, to successively adjust the NaCl concentration to 1.48, 1, 0.6, and 0.25 M. Nucleosome formation was then verified on a 6% native gel. The gel was pre-run for 1 hour at 100V at 4°C in 0.5x TBE (0.5 M Tris-Boric acid, and 0.5 M EDTA) and samples were electrophoresed for 1 hour at 100V. The electrophoretic mobility shift assay gels were soaked in H_2_O and 10 µl of ethidium bromide. Fractions that contained little free DNA were used, as it improves the stability of the nucleosome core particle (NCP).

### Histone methyltransferase assay

Methyltransferase assays were performed using 0.2 µM NPC incubated with increasing amounts of SETD2 or NSD1 in 6 µl of KMT reaction buffer (50 mM Tris pH 8.5, 5 mM MgCl_2_, and 0.2 mM DTT) and 2 µl of radioactive S-adenosyl-L-methionine, with the volume adjusted to 30 µl with mqH_2_O. The reactions were allowed to proceed overnight at room temperature and were stopped by adding 7 µl of 5x sample loading buffer and by boiling the samples for 5 minutes. The samples were resolved on a 16.5% Tris-Tricine SDS-PAGE gel using a 100 mM Tris, 100 mM tricine, and 0.1% SDS running buffer. Gel was Coomassie stained, incubated with enlightening solution for 30 minutes, then rinsed with mqH_2_O. Gels were dried for 1 hour at 80°C, autoradiographed for 3-5 days at −80°C, and signals were analyzed using ImageJ^75^.

### DNMT3B^PWWP^ crystallization, structure determination, and Setd2 modeling

Equimolar concentration of DNMT3B^PWWP^ and a trimethylated form of K36 (K36me3) of histone H3.3 peptide corresponding to residues 21 to 46 were mixed and incubated on ice for 1 hour. Crystals of the DNMT3B^PWWP^/H3.3K36me3 (21-46) complex were obtained using various sparse matrix screens. Conditions yielding preliminary crystals were further optimized, and diffraction quality crystals were obtained in 1.6 M potassium citrate, pH 6.0, at room temperature. Crystals were harvested and cryo-protected in paratone N oil. A complete dataset was collected using a Micromax 007-HF and integrated using HKL2000. The structure of the DNMT3B^PWWP^ was solved by molecular replacement using Phaser^76^ and DNMT3B^PWWP^ (5CIY) as a search model. The model was completed with approximately 20 rounds of refinement and manual building using Phenix^77^ and Coot^78^, respectively.

The homology model of the Set2d bound to a nucleosome bearing the T45I mutation on histone H3.3 T45I was generated by using Swiss-PdbViewer with the cryo-structure of Set2d bound to the nucleosome (Protein Data Bank structure accession no. 6nzo) as a template. The model was subsequently submitted to the Swiss-Model server (http://swissmodel.expasy.org/) for refinement.

